# Symmetry-breaking in adherent pluripotent stem cell-derived developmental patterns

**DOI:** 10.1101/2022.12.20.521167

**Authors:** Daniel Aguilar-Hidalgo, Joel Ostblom, M Mona Siu, Benjamin McMaster, Tiam Heydari, Nicolas Werschler, Mukul Tewary, Peter Zandstra

## Abstract

The emergence of the anterior-posterior body axis during early gastrulation constitutes a symmetry-breaking event, which is key to the development of bilateral organisms, and its mechanism remains poorly understood. Two-dimensional gastruloids constitute a simple and robust framework to study early developmental events in vitro. Although spontaneous symmetry breaking has been observed in three dimensional (3D) gastruloids, the mechanisms behind this phenomenon are poorly understood. We thus set out to explore whether a controllable 2D system could be used to reveal the mechanisms behind the emergence of asymmetry in patterned cellular structures. We first computationally simulated the emergence of organization in micro-patterned mouse pluripotent stem cell (mPSC) colonies using a Turing-like activator-repressor model with activator-concentration-dependent flux boundary condition at the colony edge. This approach allows the self-organization of the boundary conditions, which results in a larger variety of patterns than previously observed. We found that this model recapitulated previous results of centro-symmetric patterns in large colonies, and also that in simulated small colony sizes, patterns with spontaneous asymmetries emerged. Model analysis revealed reciprocal effects between diffusion and size of the colony, with model-predicted asymmetries in small pattern sizes being dominated by diffusion, and centro-symmetric patterns being size-dominated. To test these predictions, we performed experiments on micro-patterned mPSC colonies of different sizes stimulated with Bone Morphogenetic Protein 4 (BMP4), and used Brachyury (BRA)-GFP expressing cells as pattern readout. We found that while large colonies showed centro-symmetric BRA patterns, the probability of colony polarization increased with decreasing sizes, with a maximum polarization frequency of 35% at ∼200μm. These results indicate that a simple molecular activator-repressor system can provide cells with collective features capable of initiating a body-axes plan, and constitute a theoretical foundation for the engineering of asymmetry in developmental systems.

## Introduction

During early mammalian embryogenesis, gastrulation gives rise to the formation of the three germ layers, from which every cell type in the body will develop. It is also during gastrulation that the second body axis is laid down orthogonal to the already established proximal-distal axis. The emergence of this anterior-posterior (AP) body axis breaks the rotational symmetry in the epiblast, and allows for the organization of cell populations into polarized patterns. This symmetry-breaking event is key to development of bilateral organisms, and its mechanism remains poorly understood.

Most of our understanding about the processes that govern the emergence of the AP body axis come from *in vivo* studies of embryos. However, the complex developmental context of the embryo makes it difficult to disentangle and isolate individual mechanisms. To work around this technical limitation, the use of pluripotent stem cells (PSC) as embryo-like systems *in vitro* to study early developmental events has been used as a model system for to understand the body axis formation during development (Morgani et al. 2018; Amadei et al. 2022; Harrison et al. 2017). It has been observed that three-dimensional (3D) aggregates consisting only of PSCs occasionally express the primitive streak (PS)-marker BRA in a polarized manner (Harrison et al. 2017). Tight control over the number of cells per aggregate (∼300) provides robust single region localization of BRA and CDX2 (a marker of posterior embryonic regions) expression as well as elongation of the aggregates (Turner et al. 2017). Endogenous signaling, such as the WNT and NODAL morphogen pathways, is integral to the induction of BRA in mouse pluripotent stem cells (mPSCs) (Morgani et al. 2018; Turner et al. 2017)and exogenous supplementation of activators for these pathways has been shown to enhance both patterning localization reproducibility and shape elongation (Turner et al. 2017), including axial organization of gene expression (Moris et al. 2020). These elongated aggregates lack anterior structures, and the expression localization is somewhat reminiscent to what is seen in the caudal regions of the embryo at the onset of PS formation and indicates the potential presence of an anterior-posterior axis in the aggregates.

An important outstanding question is how polarized fate induction occurs among PSCs lacking the instructional extraembryonic regions. Micropatterned PSC culture systems offer a systematic and robust platform where to address this question. Results from micropatterning studies so far indicate that cell aggregates grown adherent to a substrate (adherent systems) are only capable of symmetric fate induction (Warmflash et al. 2014; Tewary et al. 2017; 2019; Chhabra et al. 2019; Kaul et al. 2022), even when micropatterned on asymmetric shapes (Heemskerk 2019; Ostblom et al. 2019; Muncie et al. 2020), where the equivalent of the posterior end of the embryo faces outwards and the anterior end faces inwards. Polarized fate induction patterns in adherent systems have only been observed when the induction signal is presented asymmetrically (Manfrin et al. 2019). However, it has been shown in intestinal organoids that the induction pattern is sensitive to the confined geometry of the cellular aggregate, leading to polarized patterns as a result of differences in the local cell packing and morphology (Gjorevski et al. 2022), suggesting that specific sizes and shapes may modulate the location of induction signals within an otherwise symmetrical aggregate.

Hitherto, the studies on PSC aggregates suggest that adherent systems might lack an inherent property present in non-adherent aggregates that is necessary for spontaneous asymmetric cell fate induction, or that further study on the relationship between induction pattern and micro-patterned geometry is needed. Ishihara and Tanaka propose that tissue engineering applications require robust results as shown in symmetric 2D adherent systems, and postulate a connection between the so-called robustness of the experimental platform, and the capability to achieve symmetry breaking (Ishihara and Tanaka 2018).

Here, we demonstrate that symmetry-breaking in 2D adherent stem cell colonies can be predictably specified as a function of the size of the micropatterned area relative to the length-scale of the morphogenetic signal, suggesting that polarization in patterns is dependent on the self-organization of activating signals within a given cellular domain. A systematic experimental analysis led us to identify regimes where polarization of the BRA domain in mPSC micropatterned colonies could be predicted, and determine dependencies between induction pattern, colony size and cell density. We also show that the dynamic emergence of both polarized and centro-symmetric patterns can be explained by the same molecular Turing-like mechanism that localizes signaling concentration gradients relative to the size of the micropatterned area. Collectively, these results suggest a developmental coupling between the activity range of morphogenetic signals and the size of the embryo for the formation of the body plan. This work open the door to engineering strategies for the control of asymmetric patterns in organoids.

## Model Development and Behavior

### Self-organized reaction-diffusion model predicts spontaneous symmetry-breaking patterns as a function of the colony size and morphogen dynamics

To guide our understanding of developmental pattern formation, and to enable the *in-silico* exploration of the effects of key parameters such as pattern size and morphogen concentration on the shape and organization of micropatterned colonies, we developed a reaction-diffusion model where the transport dynamics of one or more diffusible molecules permits the formation of long-range concentration profiles that form structured signaling patterns with specific length-scales (Müller et al. 2013; Aguilar-Hidalgo et al. 2018). The resulting shape of the signaling pattern in these models depends on the relative size of the spatial domain over which these molecules diffuse, and the boundary conditions of the system.

If these diffusible molecules couple in a Turing-like Activator (A)-Repressor (B) form, such that A activates the production of B, and B limits the production of A, concentration profiles of these molecules can self-organize within their spatial domain (Turing 1952). Although these models can simulate conditions for spontaneous symmetry breaking (Ishihara and Tanaka 2018; Sozen, Cornwall-Scoones, and Zernicka-Goetz 2021), they have so far only been used to explain the formation of the observed centro-symmetrical patterns in *in-vitro* adherent PSC colonies (Tewary et al. 2017; 2019; Brassard and Lutolf 2019; Fattah et al. 2021; Kaul et al. 2022) and to mimic polarized patterns under asymmetric induction conditions in micro-fluidic systems (Manfrin et al. 2019).

Importantly, the validity to the application of these models to 2D adherent PSC cultures has been challenged by the availability of activating signal in the media from the apical direction (Etoc et al. 2016; Chhabra et al. 2019). It has been argued that isotropic availability of the activating signal would provide homogeneous patterning in the colony, as the apical boundary of the colony should respect the same fixed boundary conditions imposed at the colony edges and the intrinsic length-scale of the concentration gradient, which is typically much larger than the height of the colony. A common characteristic in the models applied to PSC studies is that boundary conditions are fixed to either a constant concentration or flux values for A and B (Tewary et al. 2017; 2019; Kaul et al. 2022), which can constrain the number of solutions or induction patterns that they can undergo. Based on these observations and inputs, we hypothesized that changing the boundary conditions such that they can respond to the self-organization of the patterning within the colony could allow for the observation of spontaneous symmetry-breaking events in simulated micropatterned PSC colonies.

We thus developed a minimal 2-component reaction-diffusion system of the Turing-like activator-repressor type (Turing 1952; Werner et al. 2015) with reactive boundary conditions (Dillon, Maini, and Othmer 1994; Erban and Chapman 2007), where we represent the influence of the activating signal by applying signal-intensity-dependent flux at the colony boundary (see **Eqs. 1-4**). Under certain parametric conditions, this model can self-organize concentration patterns of the activator and repressor molecules. Interestingly, the correlation between the signal intensity and the flux at the boundary translates the self-organization of these patterns into dynamic changes in the boundary conditions, which allows for a broader spectrum of steady-state solutions than what was observed in models with constant boundary conditions.

This innovation in the boundary conditions provides a mechanistic basis to the temporal and spatial inhomogeneities in the cellular Bone Morphogenetic Protein 4 (BMP4) uptake that have previously been empirically imposed (Etoc et al. 2016). This interpretation of the boundary flux allows for both symmetry breaking to occur and satisfies previous concerns about the formation of centro-symmetrical patterns when considering flux coming to the cells from the apical direction (Etoc et al. 2016; Chhabra et al. 2019). This is possible since flux at the boundary will adapt to the self-organization of the intensity of the morphogenetic signals (**Fig. 1**).

**Figure 1:**
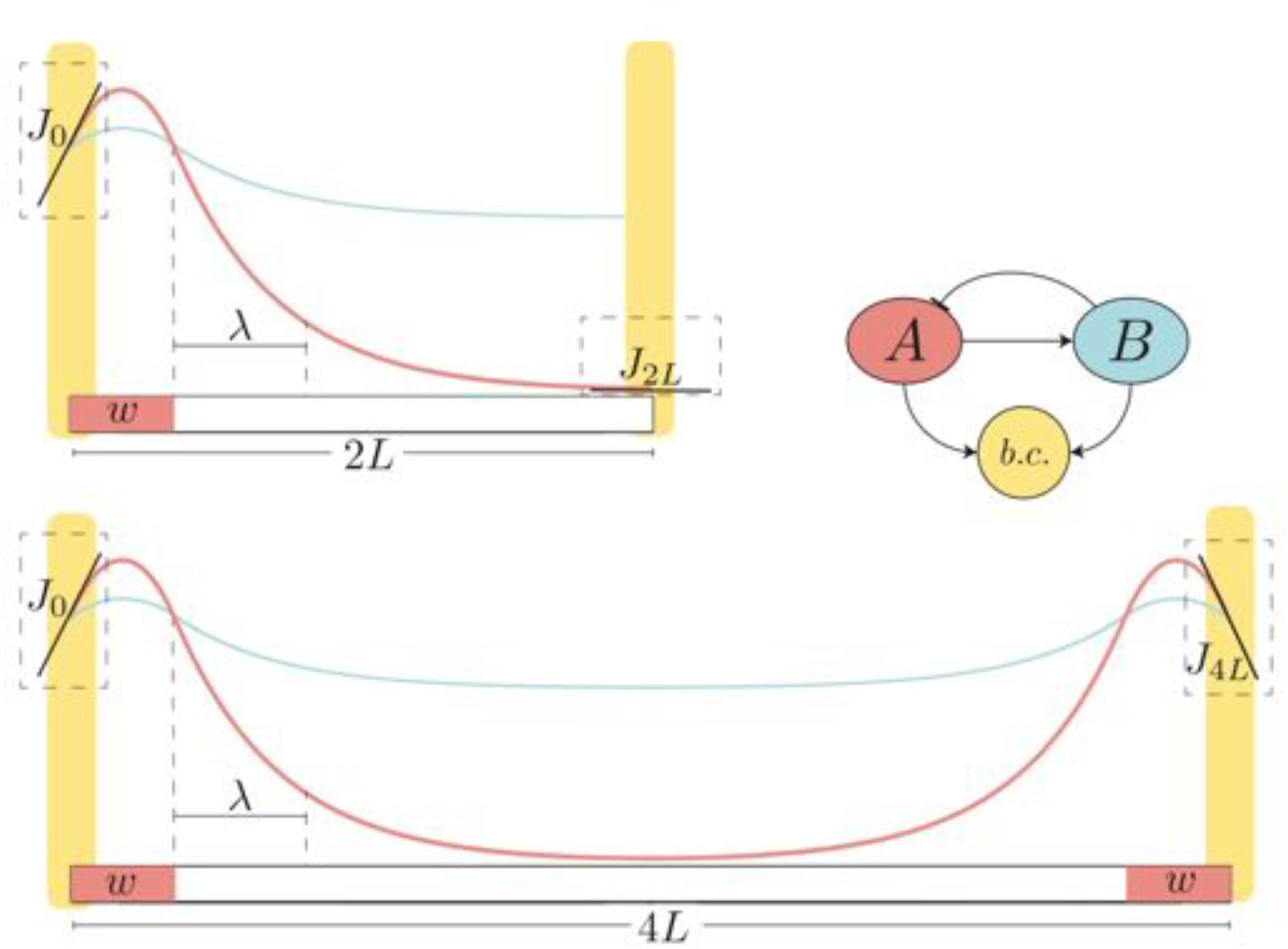
Scheme of our Turing model with self-organized boundaries. Scheme of a one-dimensional domain of size *2L* (top) and *4L* (bottom), where two diffusible molecules *A* and *B* self-organize concentration gradients following the sketched interaction motif (right). Molecules *A* and *B* are produced and secreted in the region where the concentration of A is larger than the concentration of *B*, with production function *P(A,B)=A*^*h*^*/(A*^*h*^*+B*^*h*^) (for a large value of *h*, the production function turns into a step function defining a source region of width *w*), see **Eqs. 1**,**2**. This mechanism self-organizes concentration gradients of *A* and *B* given that the concentration gradient of *B* has a length-scale much larger than the length-scale λ of *A*, which is defined as the distance from the source of *A* at which the concentration of *A* has decreased in *1/e*. This self-organized system will accommodate concentration peaks of *A* within the size of the domain depending on the ratio *L/*λ. To allow this system to show symmetry-breaking events, we linked the flux *J* at the edges of the spatial domain proportional to the concentration of *A* and *B*, see **Eqs. 3**,**4**. Thus, for a small value of *L/*λ, the concentration gradient of *A* can develop a peak in one side of the spatial domain, leading to a polarized pattern (top). Within this framework, enlarging the spatial domain for a constant λ will result in the accommodation of a second intensity peak in a centro-symmetric form (bottom).

Mechanistically speaking, our self-organized system will try to accommodate periodic patterns of high and low activator concentration spatially distributed within the colony area as a function of the wavelength (length-scale) of this periodic pattern, and the boundary conditions. According to this, a large length-scale with respect to colony size, will only be able to accommodate one high concentration peak, while a small length-scale with respect to the colony size may be able to accommodate multiple high and low concentration peaks (**Fig. 1**). In our circular colonies and for a constant activator length-scale, this translates to one peak, either in the colony center or edge, for small colony sizes. Larger size colonies with the same length-scale contain multiple high-concentration peaks that are centro-symmetrically distributed (**Fig. 2A**). This patterns can be shaped as ring-like patterns, spotted patterns, or a combination of both (**Fig. 2B**).

**Figure 2:**
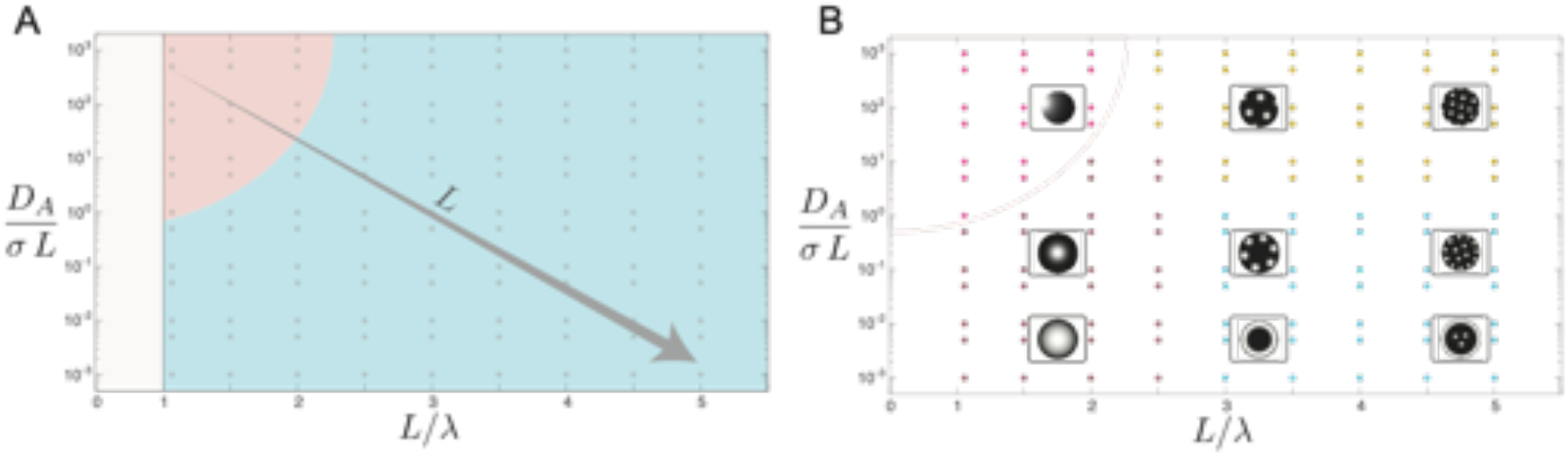
Phase diagram for pattern symmetry. Phase diagram showing solutions of the non-dimensional model **Eqs. 5-8** for values of parameters *δ* = *D*_*A*_/(*L σ*) and Λ = *L*/*λ* (dots). A) Solutions for high δ and low λ show symmetry-breaking in the concentration pattern of the activator A (pink region). Blue region shows solutions with centro-symmetric patterns. Note that an increase of the colony radius L implies a trajectory from symmetry-breaking to centro-symmetry. Note that patterns are not stable for *L*/*λ* ≤ 1 (gray region). B) Classification of solutions by pattern shape shown in colored dots with representative group image: Polarized (pink), dotted (yellow), dome (brown), ring-like distribution (cyan). Note that larger relative colony size allows for the emergence of multiple concentration peaks distributed interior to the colony, as observed in (Tewary et al. 2017; 2019). Additionally, as shown in (A), an increase in the colony radius L provides a trajectory from polarized patterns at small colony size to ring-like patterns at large colony size. Parameters: *D*_*B*_/*D*_*A*_ = 30, *k*_*B*_/*k*_*A*_ = 2, *v*_*B*_/*v*_*A*_ = 4 (Werner et al. 2015).

Regardless of colony size, the pattern formation mechanism remains the same, and with constant parameters, the length-scale of the concentration gradient and the time-scale of the pattern formation are independent on the colony size. However, the overall dynamics of the gradient formation depends on the pattern shape and the initial conditions. From a stability point of view, a centered peak in small colonies is an unstable fix-point for zero or low activator flux through the colony edge (Werner et al. 2015). Thus, starting the pattern formation dynamics from homogeneous concentration conditions subject to noise, results in a concentration peak emerging within the colony area and moving towards the closest edge location to develop a concentration gradient as a stable fixed point. The same mechanism applies in larger colonies. In this case, more than one high concentration peak can develop. Radial symmetry results in stabilization in a centro-symmetric manner leading to broken and full ring-like patterns, (see **Fig. 2**).

The equations of this model read:

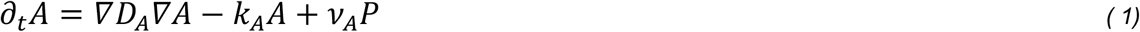

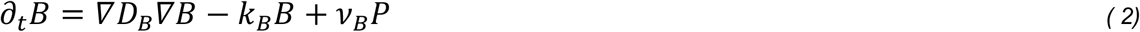

where *∂*_*t*_ denotes derivative with respect to time, ∇ is the gradient operator, *D*_*i*_ is an effective diffusion coefficient, *k*_*i*_ is an effective degradation rate and *v*_*i*_ is a production rate for *i* = *A, B*. The production function *P*(*A, B*) = *A*^*h*^/(*A*^*h*^ + *B*^*h*^) couples the concentration *A* and *B* in the form of a saturating function for *A*. For the sake of simplicity, we consider *h* to be a large number such that we approximate *P*(*A, B*) as a step function, where *P* = 1 for *A* > *B*, and *P* = 0 otherwise (Werner et al. 2015). Note that *D*_*i*_ and *k*_*i*_ are effective parameters that may encompass the effect of elementary transport events such as free diffusion, internalization, recycling, and degradation (Aguilar-Hidalgo et al. 2019). With boundary conditions:

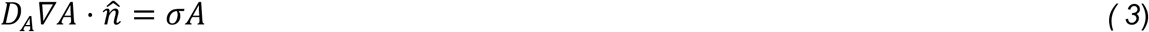

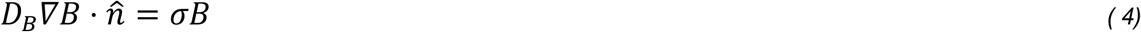

which define the diffusive flux, *J=D*_*A*_∇*A*, of *A* and *B* through the boundary, in its normal direction 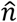, as linearly proportional to the concentration of *A* and *B* at the boundary with proportionality constant *σ*. Note that the flux through the boundaries changes dynamically and responds to the self-organization of A and B.

To simplify the analysis of these equations we choose to transform them to non-dimensional equations and consider one spatial dimension *x*. We then define a non-dimensional spatial coordinate *r=x/L*, with *L* the radius of the simulated micropattern, and non-dimensional time *τ* = *k*_*A*_*t*. This leads to the following non-dimensional form of our equations:

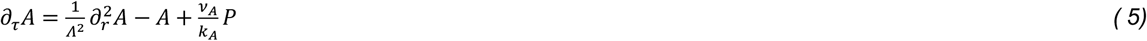

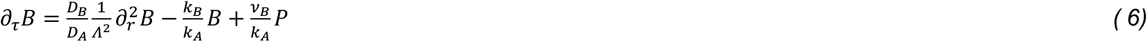

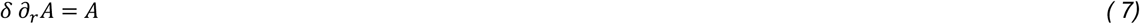

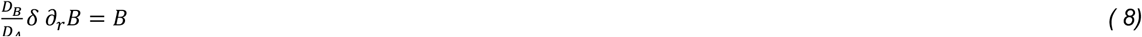

Where non-dimensional colony radius Λ = *L*/*λ* with 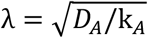 the decay length of the graded intensity profile of *A*, and non-dimensional diffusive flux coefficient 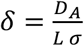. We first solved our model for steady-state in one non-dimensional spatial dimension (see sup. Mat). Note that these solutions are transferable to higher dimensions. The solution for the activator equation in a non-dimensional form is a function of the relative colony radius Λ = *L*/*λ*, see eq. (5) for the solution kernel where 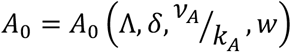 a constant pre-factor.

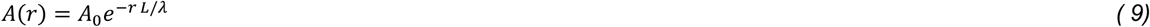

We next ran simulations sweeping free parameters Λ and *δ* to explore the space of patterns. As initial conditions for these simulations, we chose homogeneous intensities subject to noise to ensure the simulation lands in a stable fix point. For non-dimensional ratios *D*_*B*_/*D*_*A*_, *k*_*B*_/*k*_*A*_, *v*_*A*_/*k*_*A*_ and *v*_*B*_/*k*_*B*_ that ensure Turing instability (see Sup. Mat. and (Werner et al. 2015)), parameters Λ and *δ* define the phase space of pattern solutions. Note that these two parameters show a countereffect between system size *L* and diffusion *D*_*A*_ such that increasing *L* increases Λ and decreases *δ*, while increasing *D*_*A*_ decreases Λ and increases *δ*. This countereffect creates two distinct pattern regions where for high *δ* values and low Λ values, the pattern is diffusion dominated, and for low *δ* values and high Λ values, the pattern is size dominated (**Fig. 2A**). We find that in the diffusion dominated region patterns undergo spontaneous symmetry breaking, while the size dominated region shows centro-symmetric patterns. In the latter region, low Λ values show dome-like patterns with increasing width as *δ* decreases, which implies a shorter flux range from the colony edge. For high Λ values, patterns are size dominated and while for high flux patterns scatter peaks throughout the colony, they gradually organized closer to the edge when lowering *δ* to eventually form solid structures along the colony edge in the form of ring-like patterns (**Fig. 2B**). Interestingly, for constant parameters and scanning the micropattern radius *L*, our system naturally transitions between polarized patterns for small micropatterns to ring-like patterns for large micropatterns following the left-to-right diagonal in our phase diagram (**Fig. 2A**)

Our centro-symmetric solutions agree with observations previously reported in hPSC colonies (Tewary et al. 2017; 2019), and suggest that the conditions set in those experiments allows for a size dominance in the patterning where the BMP4 diffusive range is very restricted to the colony edge. This is consistent with the emergent empirical interpretation of high BMP4 receptor density (and thus high binding flux) at the edge of the colony, as proposed by Etoc et al. (Etoc et al. 2016). Additional concentration peaks, as previously observed (Tewary et al. 2017; 2019), would emerge as a result of the self-organization of periodic concentration patterns due to the relative increase in the colony size with respect to the length-scale of the signaling gradient Λ = *L*/*λ*.

Our observation of spontaneous symmetry breaking at small micropattern size, leading to polarized patterns suggest that transport through the colony is dominated by diffusion, and not necessarily restricted to the colony edge, where a small local perturbation within the colony will self-organized the accommodation of one concentration peaks of *A* towards one side of the colony edge. These results suggest that there might be experimental conditions where polarized and centro-symmetric patterns can emerge as a function of the size of the colony relative to the length-scale of the morphogenetic signal. This pattern formation mechanism predicts that polarized patterns may be found in small colonies relative to the morphogen length-scale, and could explain both centro-symmetric and polarized patterns, with the latter emerging from spontaneous symmetry breaking events.

Further, the morphogen length-scale 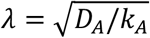 depends on the morphogen transport properties, which can be cell-density (receptor binding) dependent (Müller et al. 2013; Aguilar-Hidalgo et al. 2019). Cell density may impact the sensitivity of morphogen absorption, which refers to our model effective parameter *σ*, as a result of hindered transport (e.g. proximity to cell membrane (Eloul and Compton 2016; Chio and Tse 2020), changes in effective receptor density, which results in changes in ligand-receptor binding and internalization rates (Aguilar-Hidalgo et al. 2019). Additionally, higher cell densities may reduce the effective diffusion coefficient *D*_*A*_ as denser cell-packing could hinder molecular transport. The countereffect of parameters *σ* and *D*_*A*_ in our non-dimensional parameter *δ* show that modifying cell density can either increase, decrease or keep constant *δ* depending on their relative effect on this parameter. The effect in our parameter Λ is clearer as a decrease in *D*_*A*_ will result in a smaller *λ* and thus a shorter concentration gradient, which will increase Λ.

In the next section, we explore the effect of cell density and colony size in the Bra pattern distribution. Note we consider Bra expression as a reporter of the BMP4 signal (Bernardo et al. 2011). To find a correlate between cell density and *λ*, we set out to scan a wide range of seeding cell densities with the aim of finding an interval that could provide polarized and centro-symmetric patterns. For colony size, we scanned a colony diameter range that could potentially show both polarized and centro-symmetric patterns, as predicted in **Fig. 2B**. Considering a constant value for *λ*, polarized patterns should emerge in colonies 2-3 times smaller than the colonies where centro-symmetric ring-like patterns are observed (diameter of 600-700μm) (Tewary et al. 2017; Morgani et al. 2018). To be conservative, we will scan colony diameters between 100μm-800μm (*L*=50-400μm).

## Results

### Polarized Bra patterns emerge as a function of seeding cell density and colony size as control parameters

In the Model Development section above, we introduced an Activator-Repressor system, which predicts that polarized pattern would emerge in small circular micropatterned colonies, while centro-symmetric ring-like patterns would emerge in large colonies. We further discussed that the likelihood of developing such polarized patterns may be sensitive to variations in cell density. We and others (Tewary et al. 2017; Morgani et al. 2018) have observed Bra+ cells organized in ring-like patterns for colony diameters larger than 600μm. We sought to explore if reducing the diameter to 200μm would promote polarization in cellular patterns. This is coincident with both the diameter range where our model suggests that polarization might occur, and with the size of the epiblast in the mouse embryo (Sozen, Cornwall-Scoones, and Zernicka-Goetz 2021; Orietti et al. 2021). To explore the effect on patterning of seeding cell density, we doubled density in the interval 10k to 80k to find optimal values that may allow for the emergence of polarized patterns. Mouse PSCs were primed for 4 days in N2B27 media supplemented with FGF, Activin A, and KOSR (NBFAK) before differentiating in N2B27 media supplemented with FGF, Activin A, Wnt, and BMP4 (NBFAWB) for 48 h in the 96-well plate. We noticed striking differences in both the extent of marker expression and its spatial organization (**Fig. 3A**) with changes in seeding density, where the proportion of the Bra+Sox2-region increased with lower seeding.

**Figure 3:**
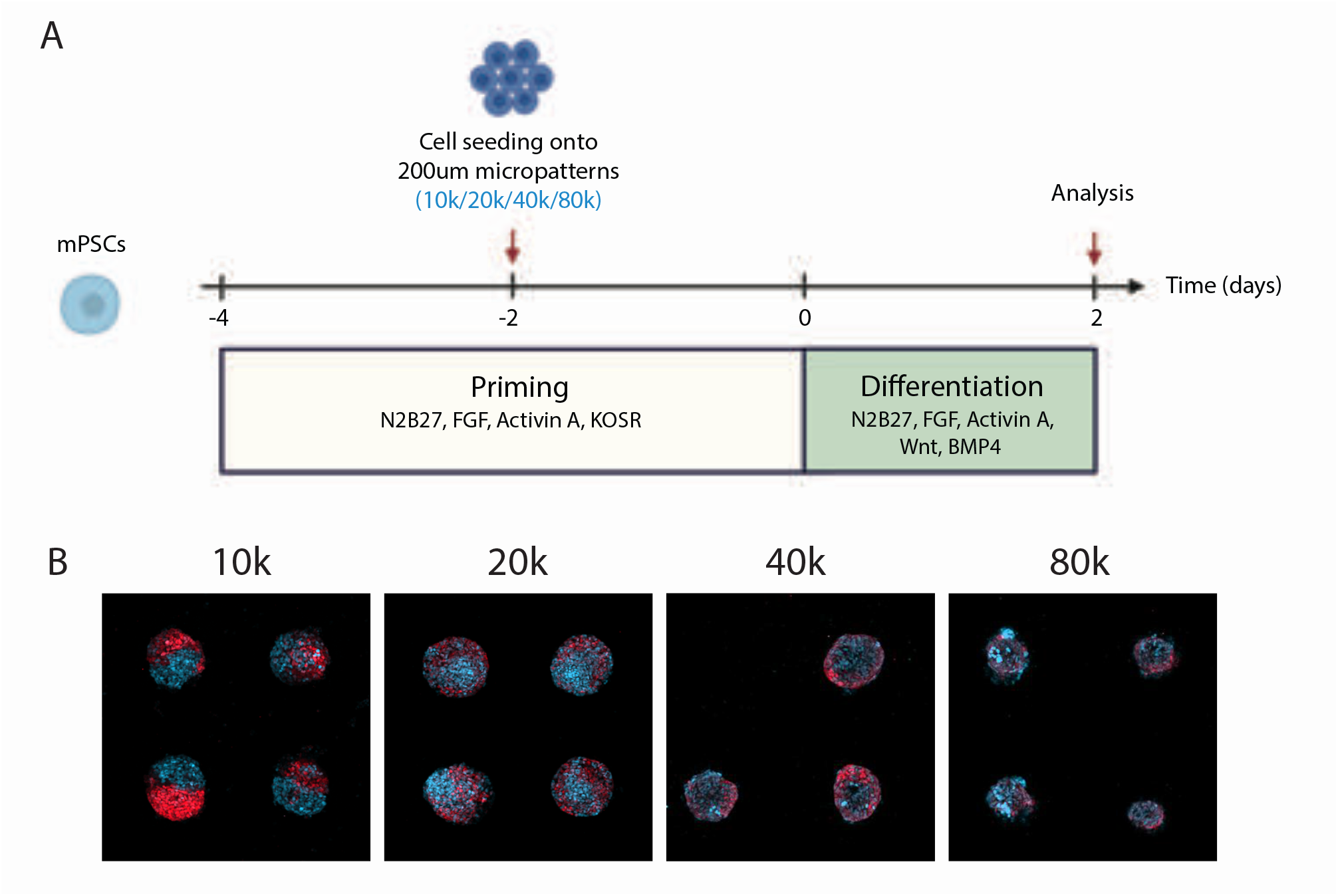
A critical range of cell densities support polarized expression of Bra and Sox2. **A)** Schematic of the experimental outline. mPSCs were primed in N2B27 with FGF, ACTIVIN A, and knockout serum replacement (KOSR) for 48h before plating onto 200 μm micropatterns at different densities. Micropatterned colonies underwent priming for another 48h and were then differentiated in N2B27 with FGF, Activin A, Wnt, and BMP4 for 48h. B) Representative microscopy images of colonies 200 μm in diameter with seeding density of 10k, 20k, 40k, or 80k. Bra is shown in red and Sox2 in cyan. The side of each image is 750 μm.

Interestingly, in colonies of 200 μm diameter Bra+ and Sox+ regions rarely organize as a centered cluster of Sox2+ cells with a surrounding region of Bra+ cells along the edge. Rather, regions positive for either Bra or Sox2 are often offset from the colony centroid and localize at different poles of the colonies. We observed that this polarization was more pronounced in the lower edge of the scanned cell density range.

We now sought to explore the influence of the colony size on the pattern outcome. As our model predicts a transition from polarized to centro-symmetric patterns as a result of increasing colony size, we choose to scan cell diameter between 100 – 800 μm for a cell density of 10k cells/well, which provides clear polarized patterns (**Fig. 3B**). Thus, we primed mPSCs for 2-days after transferring them to 96-well plates containing micropatterns of 100 - 800 μm in diameter in separate wells. These cells were differentiated in N2B27 media supplemented with FGF Activin A, Wnt and BMP4 (NBFAWB) for 48 h and stained for Sox2 and Bra, see Methods for details. The smallest colonies (100 μm diameter) showed high variation in spatial organization of markers with some colonies being entirely Sox2+, some entirely Bra+, and some expressing both markers, but at opposite ends of the colony (**Fig. 4, Sup. Fig. 1**). Similar variation has previously only been reported for much smaller colonies of hPSCs, where “micro-colonies” consisting of no more than eight cells undergo all or nothing responses to inducing ligands rather than patterning fates unevenly within the colony (Nemashkalo et al. 2017). In 200-300 μm colonies, we observed the polarized expression illustrated in (**Fig. 4A-B**) as predicted above, while at 400-500 μm the Sox2+ region becomes centrally located in the colony and the Bra+ region starts to spread out along the edge of the colony (**Fig. 4A-B, Sup. Fig. S1**). These organizational events were more pronounced in bigger colonies, as can be seen for the 700-800 μm colonies (**Fig. 4B**).

**Figure 4:**
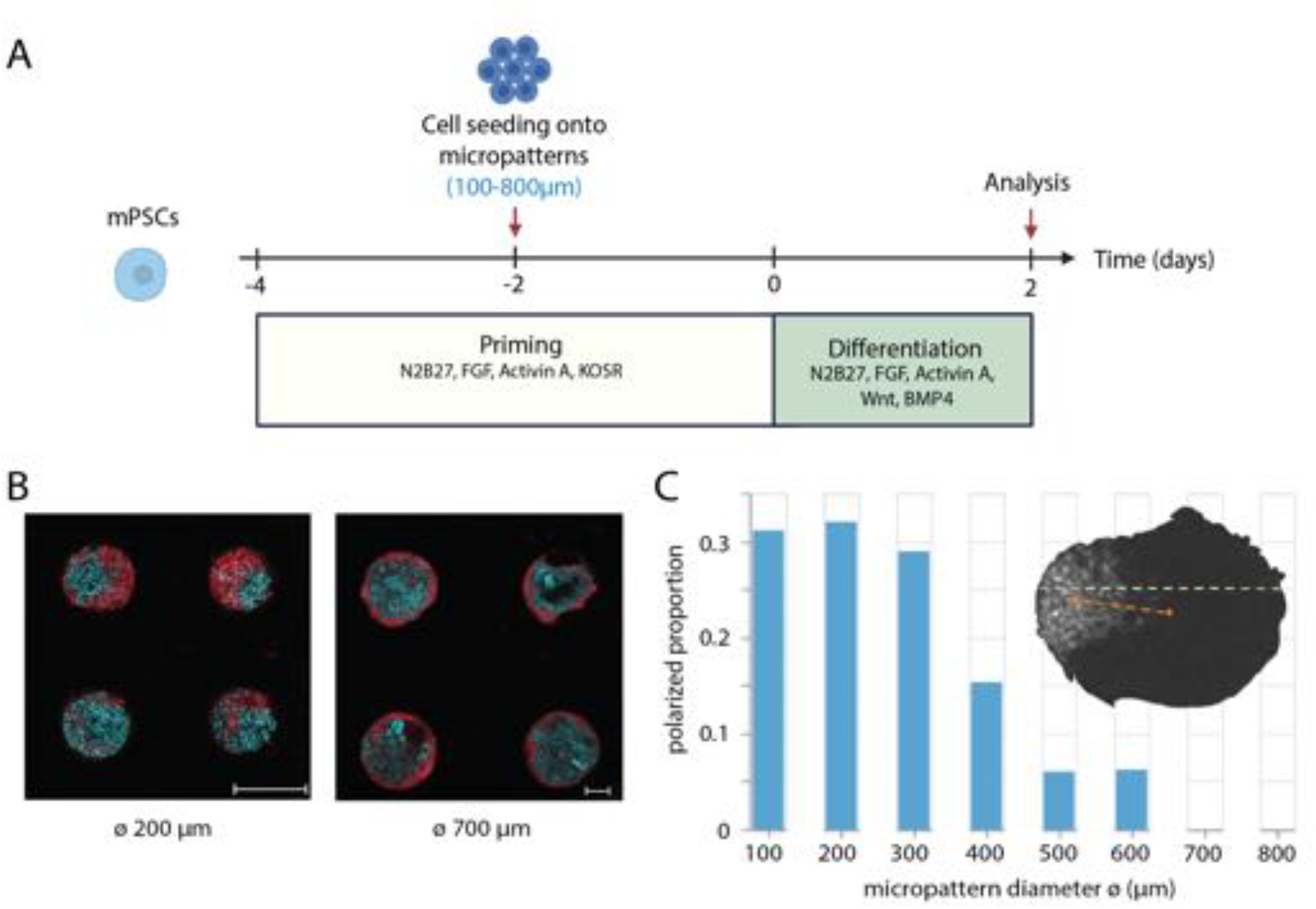
A critical range of colony diameters supports polarized expression of Bra and Sox2. **A)** Schematic of the experimental outline. mPSCs were primed in N2B27 with FGF, ACTIVIN A, and knockout serum replacement (KOSR) for 48 h before plating onto micropatterns in different sizes. Micropatterned colonies underwent priming for another 48h and were then differentiated in N2B27 with FGF, Activin A, Wnt, and BMP4 for 48h. **B**) Maximum intensity projections of confocal slices from fluorescently stained micropatterned colonies of 200μm and 700μm diameter (image scaled to fit in the figure). Cyan = Sox2, Red = Bra. Scale bar = 200um. **C)** Proportion of polarized pattern decreases per colony diameter for seeding cell density of 8k cells/well. Inset: Example of quantification of a polarized pattern by measuring the distance between the center of mass of the Bra+ pixel region and the colony centroid (red dotted line). This offset distance is normalized to the size of the colony by dividing it with half of the length of the major axis in the colony (the diameter/major axis is marked with the yellow line).

Quantifying this localized expression by measuring the distance between the colony centroid and the center of mass for positive pixels of each marker revealed that the Bra+ region is the most offset from the centroid around 200 μm colonies as predicted by the model (**Fig. 4C**). Larger colonies of around 600 – 700 μm show ring-like patterns where Bra is expressed around the edge of the colony.

Here we have shown that polarized Bra patterns can be observed in colonies of around 200 μm of diameter, and that larger colonies provide centro-symmetric patterns. According to our model, in order for this to happen, the length-scale of the signaling gradient should be colony-size invariant and smaller than the colony radius. Otherwise, the full colony would become BRA+. To explore whether this is indeed the case, we next sought to quantify the decay length of the Bra intensity profile for different colony sizes in a range of seeding cell density values where polarized patterns frequently occur.

### Decay-length of the BRA gradient slightly decreases with cell density

As mentioned above, our model considers the decay length of the activator to be invariant to changes of the colony size. Additionally, we earlier showed that cell seeding density is a sensitive parameter to obtain polarized patterns (**Figs. 3**,**4**), and in the Model Development section we described how increments in cell density can lead to the shortening of the decay-length *λ* of the signaling gradient. If this is consistent with our experimental *in vitro* system, *λ* should stay approximately constant for changes in the colony size and decrease with increasing cell seeding density. To evaluate this hypothesis, we quantified the decay-length of the Bra gradient for different system sizes and cell seeding densities, by fitting the steady-state solutions of our model to the experimental gradients (see Sup. Mat. for details). As for cell seeding densities, we chose to evaluate the range 8-14k, as low densities showed clearer polarization in patterns (**Fig. 3**). Note that while in our model *A* corresponds to a diffusible molecule, we use Bra as a reporter of the morphogen BMP4 spatial profile (Bernardo et al. 2011), and thus measure the Bra gradient decay length.

We first quantified the average Bra expression profile from a total N=3365 circular colonies with diameter of ranging between 100μm-800μm for different cell seeding densities (8k-14k cells/well), **Fig. 5** and **Sup. Fig. S2-S4**. Here we found that for a very small colony size (diameter of 100μm), the average Bra expression peaked in the colony centered with lower expression at the colony edge. Note that for colony diameters of 200-300μm, these averaged heatmaps show a ring-like pattern as the polarization occurs at random positions per colony, please refer to **Fig. 4** and **Sup. Fig. S1** to observe single colonies. Note also that for large colony diameter (700-800μm) Bra+ cells form more irregular structures though still resembling rings in individual colonies.

**Figure 5:**
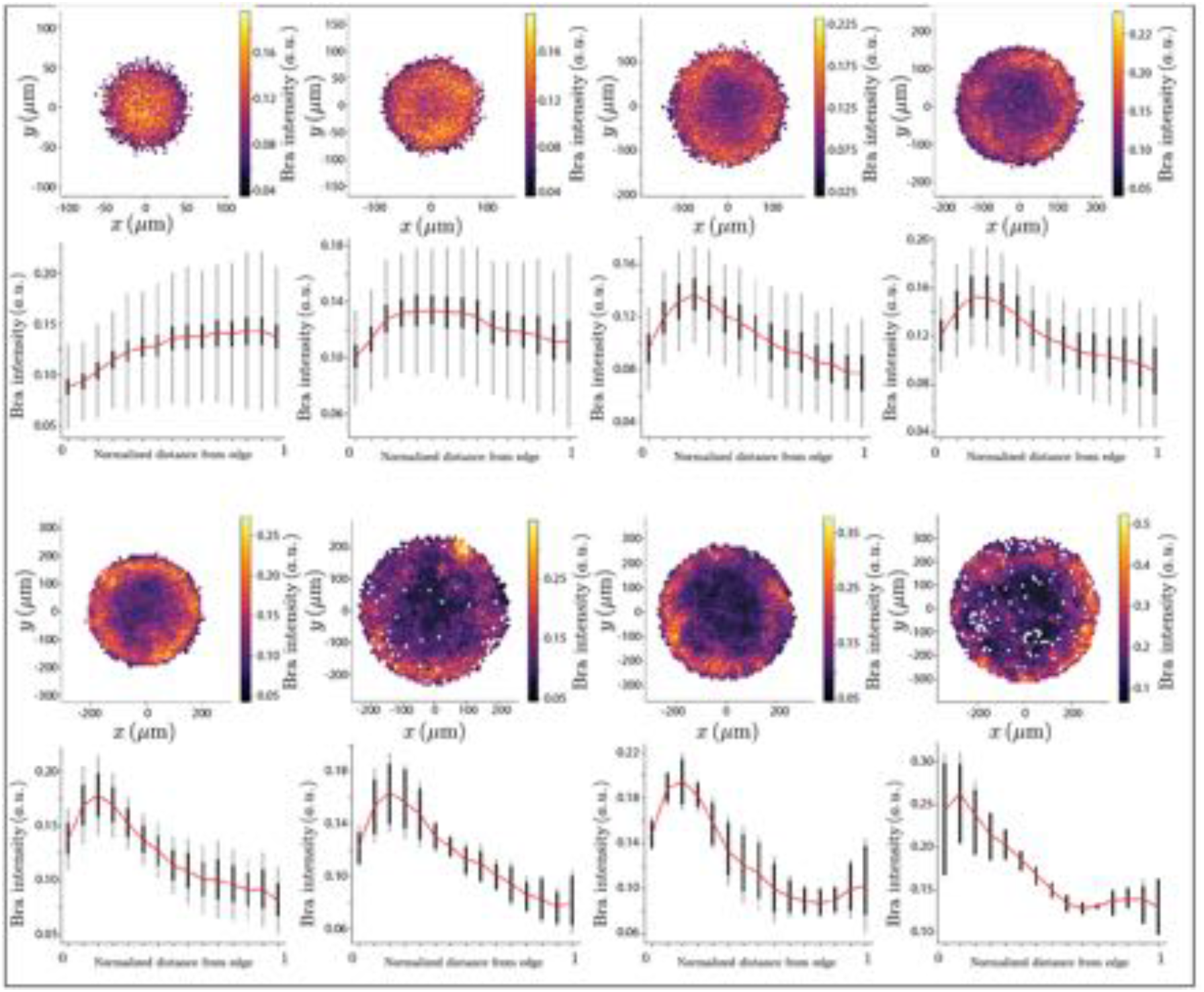
Quantification of average BRA expression gradients. Heatmaps showing the average Bra expression over a total of N=247. From upper-left corner to bottom right, the radius of the colony spans L=50-400μm. Below each heatmap, a radial quantification of Bra gradient shown in bins over a normalized radius. Red line connects the average value per bin. Thick back lines represent the 95% confidence interval for cells grouped in that bin, and light black lines the standard deviation. Cell seeding density 8k.

We then fitted the model solution in steady state to the Bra expression profiles to obtain the value for *L*/*λ* per colony size. Comparing *L* and *L*/*λ* for different colony sizes reveals an approximate linear relationship, particularly at higher density (**Fig. 6A**). A linear relationship here implies that the decay length of the Bra expression profiles, given by the slope of the linear fit, is approximately constant across colony sizes, which agrees with the model. Interestingly, the decay length appears to decrease slightly and consistently with increasing cell seeding densities, although the confidence intervals for adjacent densities are too large to make strong claims about this relationship, (see **Fig. 6B**). These results suggest that the transport of ligands within the colony may deplete with increasing cell density.

**Figure 6:**
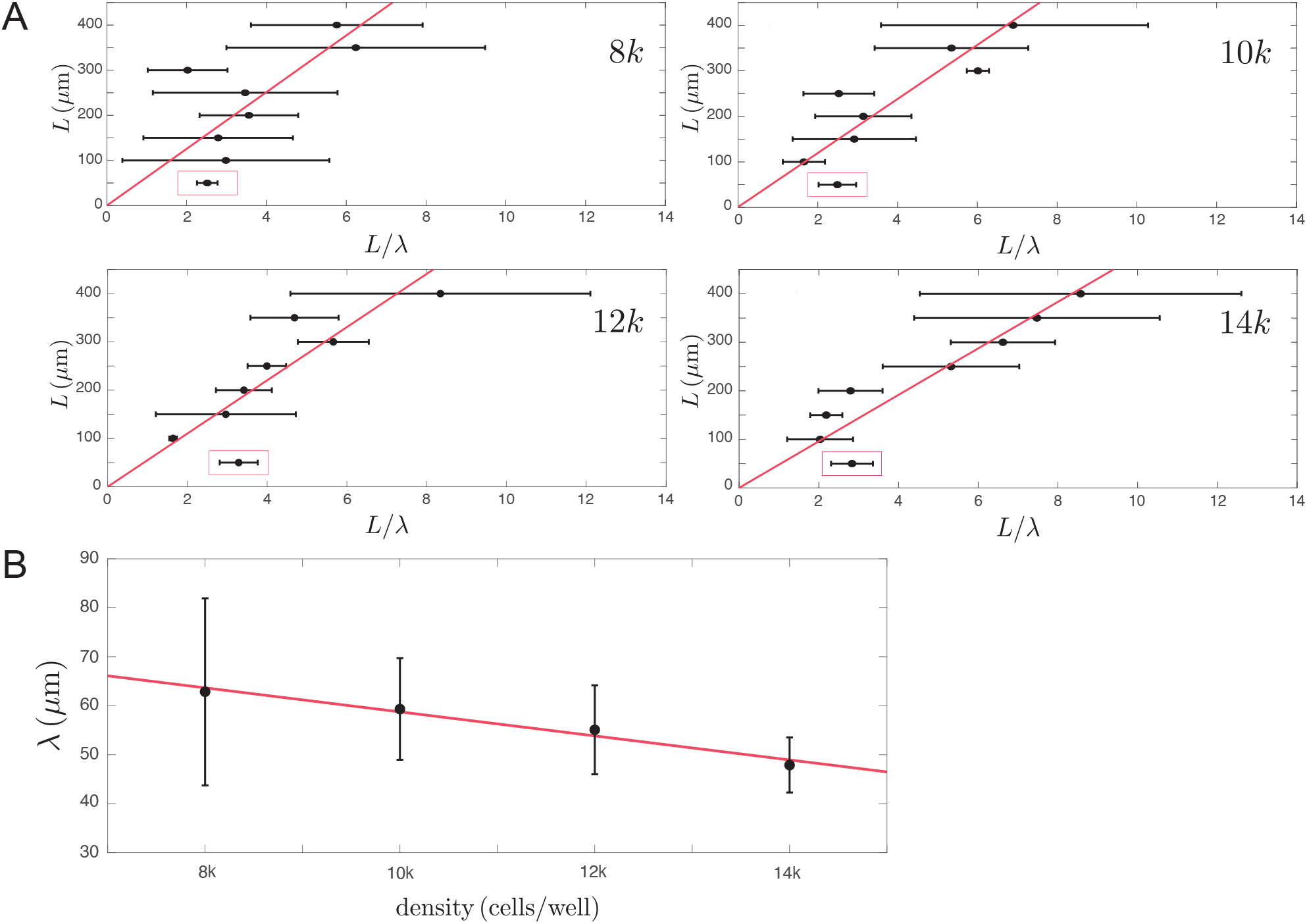
The decay length of the Bra profile slightly decreases with cell density. (A) Quantification of the decay length λ of the Bra gradient. The slope of the linear fit (red line) shows the value λ at a cell seeding density of (8k) cells per well (λ = (62.8±19.0)μm. Goodness of fit: R^2^=0.39), (10k) cells per well (λ = (59.4±10.4)μm. Goodness of fit: R^2^=0.78), (12k) cells per well (λ = (55.1±9.1)μm. Goodness of fit: R^2^=0.81), (14k) cells per well (λ = (48.0±5.6)μm. Goodness of fit: R^2^=0.90). Data in red box are excluded from the fit. (B) Variation of the decay length of the Bra gradient *λ* with the seeding cell density. Black dots represent the average lambda value, and error bars show the 95% confidence interval. Red line shows the linear fit. *λ* = *a* · *density* + *b*, with *a* = −(2.45 ± 1.29) 10^−3^ *μm*/*cells*/*well* and *b* = 83.3 ± 14.6 *μm* Goodness of fit: *R*^2^ = 0.97. Errors in quantification correspond to the 95% confidence interval.

We next simulated the emergence of patterns for measured values of λ and scanned for the colony size range we performed experiments. **Figure 7A** shows simulation results, where from increasing colony diameter (2L) pattern shape transition from polarized (100-300um, note that at 300um a second smaller peak emerges shaping two opposite poles) to homogeneous (400um) to ring-like structures (500-800um). **Figure 7B** shows an overlay of experimental results matching simulated patterns, with Bra shown in Red and Sox2 in Cyan. Note that simulated asymmetric (not polar) pattern were not experimentally found, suggesting that either those solutions cannot be observed experimentally or that a finer discretization step may be required in the experimental parameter to let the system evolve to that shape. The samples shown here correspond to a decreasing cell density trajectory from small to large colonies (see figure caption for details). How cell-density influences the ratio *D*_*A*_/*σ*. These results suggest that the formation of both centro-symmetric and polarized patterns can be explained by a Turing mechanism with self-organized boundary flux.

**Figure 7:**
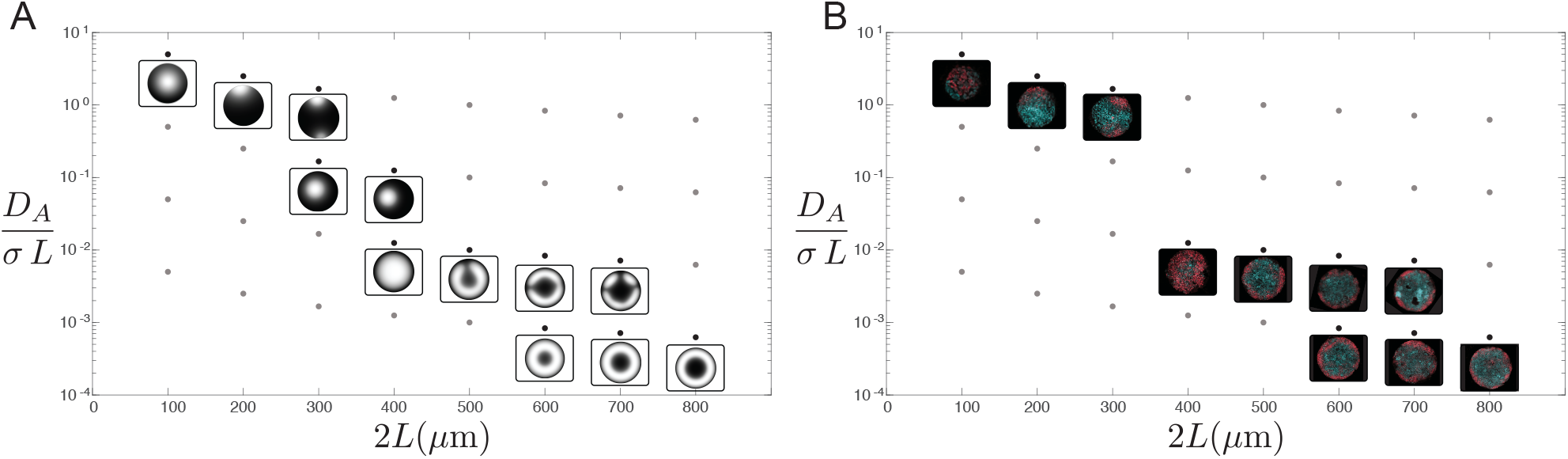
Simulation and experiment pattern shape match for measured decay length. (A) Simulation results scanning colony diameter (*2L*) and parameter σ for a fixed value of the decay length and diffusion coefficient (gray and black dots). Black dots show below simulated pattern for increasing colony diameter. (B) Overlay of experimental images that match simulation results. Parameters: *λ* = 50*μm, k*_*A*_ = 10^−4.^ *s*^−1^, *D*_*A*_ = *λ*^2^ *k*_*A*_, *D*_*B*_/*D*_*A*_ = 30, *k*_*B*_/*k*_*A*_ = 2, *v*_*B*_/*v*_*A*_ = 4, *L* = (50 − 400)*μm, σ* = (0.001 − 1)*μm*/*s*.

#### Marker polarization arises via displacement rather than induction at the poles

We next sought to investigate if the dynamics of the emergence and localization of the Bra+ region agrees with the proposed mechanism from our model where high concentration peaks self-organize as a function of the decay length of signaling molecule gradient and colony size. As mentioned in the Model Development section, this pattern formation mechanism is the same for all colony sizes. However, as a physical system, different perimeter to area ratio in the micropattern can lead to apparent differences in how the dynamics emerge while the localization of high-concentration peaks develops towards a stable region. As a reminder, our theoretical model predicted that different colony sizes will lead to different shapes of the signaling gradient (**Fig. 2**). In particular, we predicted that large colony sizes (600-800μm diameter) will show ring-like structures, while small colony sizes (200-300μm) will develop polarized patterns. The signaling gradient formation mechanism would dynamically accommodate one high concentration peak of the activating signal in the colony center or edge for small colony sizes (200-300μm), and two high concentration peaks for large colony sizes (600-700μm), with the latter centro-symmetrically distributed in a ring-like shape (**Fig. 2**).

To mimic experimental conditions, we used simulation results from **Fig. 7**, where we incorporated the experimental measure of the decay length of the Bra gradient. Observing the dynamics of the gradient formation in simulated colonies of 200μm and 600μm, we find that at early times, a concentration peak starts to form interior to the colony for both sizes (**Fig. 8A**). However, the larger colony can accommodate two concentration peaks, and by rotational symmetry they show as a ring-like structure. With time, the single concentration peak in the 200μm colony develops slightly towards the edge, while the ring-like structure in the 600μm colony consolidates as a more solid ring until they reach a steady state. In summary, we see that the same mechanism can provide a different visual cue of the pattern formation.

**Figure 8:**
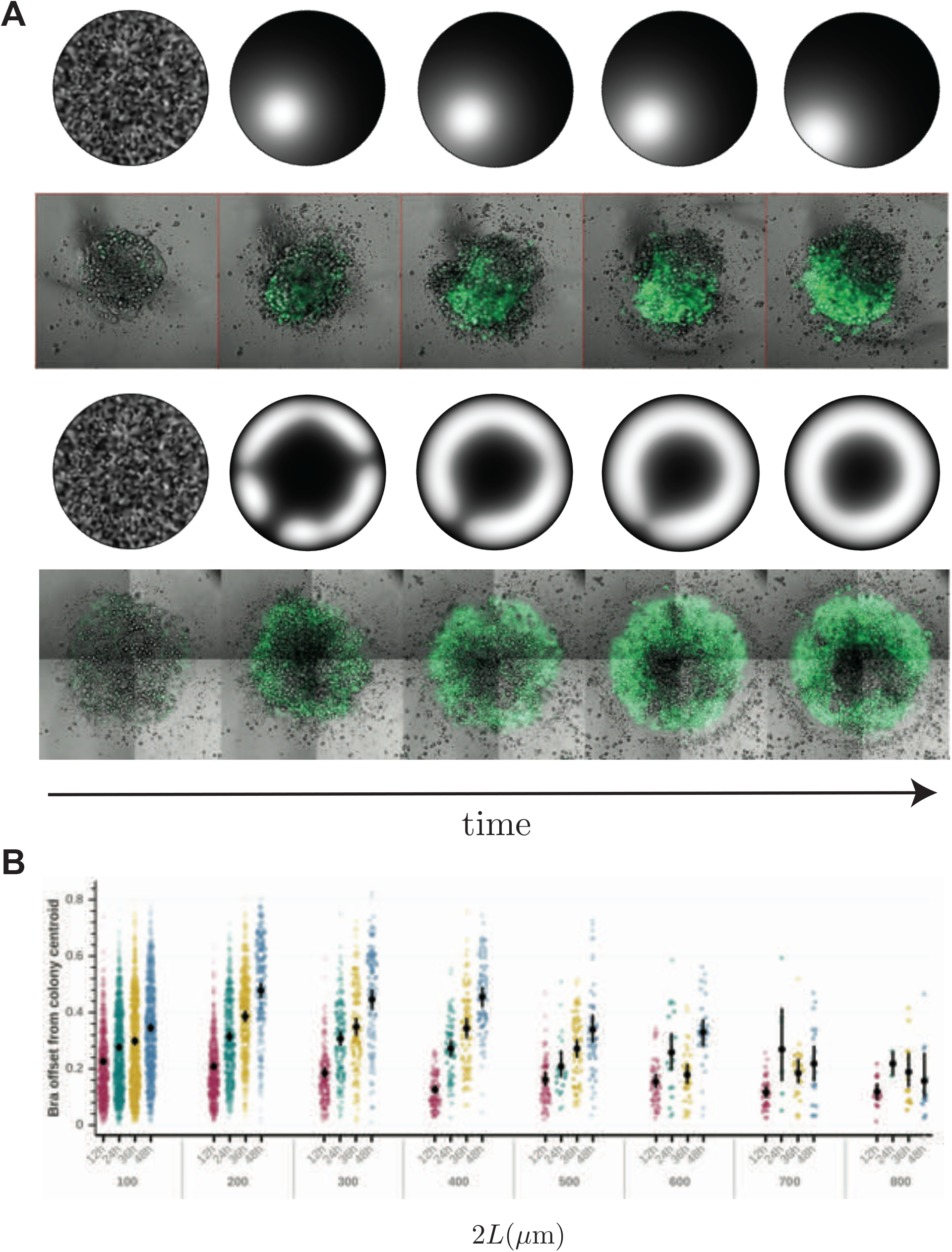
Evolution of Bra expression over time in colonies of difference sizes. (A) Representative images spaced equally in time from *in silico* and *in vitro* differentiation over 48h. The top two rows show 200 μm colonies undergoing polarized differentiation and the bottom two rows show 600 μm colonies undergoing centrosymmetric differentiation. The in vitro live imaging experiment show expression of a Bra-GFP construct overlaid with T-PMT. Simulation parameters as in **Fig. 7**. (B) Distance between the center of mass of the Bra+ region and the centroid of the colony at four timepoints during differentiation (12-48h), for colony diameter ranging between 100-800μm for seeding cell density of 8k cells/well. Distances are normalized to the colony radius, so the theoretical max value is 1 which indicates a single Bra+ pixel at the colony edge. The practical max value is just under 1 as colonies are thresholded to include only those with at least 2% Bra positive pixels to avoid artefacts. Each colored dot represents a colony positioned according to the kernel density estimate of the underlying distribution. Black dots represent averages, and error bars indicate 95% confidence intervals, so significance at p<0.01 can be inferred roughly from non-overlapping error bars and at p<0.05 from error bars that overlap by a quarter of their total length.

To investigate if the mechanism provided in our model agrees with experimental results, we followed the progression of micropatterned colonies over time by seeding cells at high and low density and assaying them at 12h intervals over two days. Intriguingly, in colony sizes <= 500

μm in diameter, the distance between the Bra+ region and the colony centroid increased gradually over time, starting already from the 12h to 24h time point (**Fig. 8A,B**). This indicates that the Bra+ region is not induced in place at the edge of the colony and then grows over time, but rather alters its location over time progressively moving away from the centroid of the colony.

These results agree with the mechanistic understanding of our pattern formation model, where polarized induction emerges interior to the colony to later move towards the colony edge, while ring-like structures centro-symmetrically close to the colony edge. This also suggests that the timescale of the induction pattern is larger than the stabilization of the signaling gradient, thus showing that the temporal progression of cell fates’ spatial organization in smaller polarized colonies occur largely through displacement after induction. This effect cannot be observed in bigger colonies where Bra+ regions are induced in largely symmetrical patterns along the edge of the colonies and do not change their location after induction.

## Discussion

In this work, we present a comprehensive analysis of the spatial organization of peri-gastrulation-like patterns in mouse stem cell adherent colonies, with particular focus on control parameters that distinguishes between centro-symmetric and polarized patterns, the latter with similarities to the organization *in vivo* of AP body axis.

While to date, polarized expression of Bra, indicative of the formation of the AP axis, has been observed only in 3-dimensional gastruloids (Turner et al. 2017; Beccari et al. 2018; Moris et al. 2020), we found that the capability to organize cell fates in a spatially polarized manner is also an inherent property of 2D micro-patterned stem cell populations, and not a unique feature of 3D cell systems. We have unlocked this feature by carefully manipulating the system size (both in terms of colony diameter and cell number). To do this systematically, we developed a simple theoretical framework based on Turing’s activator-inhibitor model to explore the patterning phase-space. We found agreement with previous observations in pattern transitions such as from ring-like patterns to a ring with internal spots (Tewary et al. 2019), and predicted conditions to transition to polarized patterns as a function of the colony size, which needs to be smaller than in the case of the emergence of ring-like patterns.

Turing models have been widely explored and the relationship between the shape of the pattern and the size of the system where the pattern forms is well understood, in particular with respect to the periodicity of the pattern length-scale within the system and its adaptation to the system’s boundary. In the stem-cell field, this type of models has been previously used to explain mechanisms of pattern formation in 2D micro-patterned systems. While these models successfully explained centro-symmetric patterns in conditions where the micropattern size permits the accommodation of more than one pattern length-scale, they have been unable to explain the formation of asymmetric patterns from homogeneous conditions which have previously also not been observed in micropatterned colonies. Moreover, the adaptation of these models in two-dimensional simulations has been controversial, as the flux of signaling molecules will be fixed and limited to the edge of the simulated micropattern, while experimentally this flux could come as well from the apical direction. Then the pattern should be completely overridden as the height of the colony is probably much smaller than the pattern length-scale. The observation of patterning in such thin micropatterned colonies, has so far been explained as a buffering from BMP4 inhibitors in the apical direction.

Here, our model applies the so-called reactive boundary conditions that defines a flux through the boundaries proportional to the concentration of the activating signaling molecule. This feature permits the flux to adapt to the self-organization of the concentration pattern within the system’s boundary. Therefore, edge regions with high concentration of the activator will show a high flux while regions with low concentration will show low flux. Interestingly, this mechanism in an experimental micropattern will explain the formation of patterns even with flux coming in the apical direction, as the flux will adapt to the self-organizing BMP4 concentration within the colony. Noteworthy, this mechanism also agrees with the observation of the relocalization of BMP4 receptors in the colony edge, as per the edge-sensing paradigm (Etoc et al. 2016), allowing a higher sensitivity to BMP4 in cells in the colony edge than in the colony center. In this sense, our model integrates the Turing’s activator-inhibitor and edge-sensing paradigms, such that edge sensing is a consequence of the dynamic self-organization of the BMP4 reception at the colony edge driven by a Turing mechanism with reactive boundary conditions.

We find that spontaneous symmetry-breaking events happen for a particular range of simulated colony sizes for fixed length-scales of the morphogenetic signals that allow for only one high-concentration peak to be accommodated within the size of the colony. This suggests that polarization of expression patterns is dependent on self-organization of activating signals within the given cellular domain. A systematic experimental analysis led us to find the predicted Bra polarization in mPSC micropatterned colonies but also to determine dependencies between induction pattern, colony size and cell density. Specifically, we found that the length-scale of the Bra intensity profile decreases with cell density.

Considering that the Bra gradient follows its upstream BMP4 gradient, this result suggests that BMP4 effective diffusion is hindered by cell packing. Transport of BMP4 throughout the colony may have a larger free extracellular component in colonies with lower cell density, while molecules may have to be transported through cells in highly dense colonies (Aguilar-Hidalgo et al. 2019). If BMP4 effective degradation rate is invariant to changes in the cell density, the timescale for the gradient formation will be independent of changes in colony size and cell density. This combined transport mechanism has already been proposed in other developmental model systems (Aguilar-Hidalgo et al. 2018; 2019; Romanova-Michaelides et al. 2022). Additionally, we observed that cell density influences the likelihood for polarization. Cell density dependency could be explained by its influence in molecular transport, which may turn into a change in the length-scale of the signaling gradient. This will effectively provide conditions for either more than one concentration peak to emerge, or to fill in the entire domain with high concentration signal, which leads to homogeneous patterning. This suggests that engineered signaling gradients could provide control of the shape of in-vitro patterning in pre-selected colony sizes and densities.

Our findings demonstrate that PSC colonies cultured in printed 2D systems contain the necessary information to undergo spontaneous symmetry breaking and elicit polarized expression of the early gastrulation marker Bra akin to its localized expression in gastrulating embryos. These results suggest extraembryonic signalling is not required for this asymmetry to occur; and that it might serve another primary purpose during development, such as stabilization of the gradient as previously speculated (Morgani et al. 2018). Our findings also support the existence of a coupling between developmentally relevant system sizes to signaling length-scales in order to give rise to cell induction and system polarization reminiscent of the formation of the AP body axis.

Our study supports the importance of extensive analysis on the coupling between cell density, system size and morphogen transport dynamics in the control of functional in-vitro tissues. Further insights in these topics could improve our understanding for how to control fate organization in cell populations and can advance both our understanding of developmental processes and how to create complex tissues with regenerative engineering.

## Methods

### Mouse embryonic stem cell culture maintenance

Cell lines used in this work include R1 (Nagy et al. 1993) and BraGFP/+ (Fehling et al. 2003) mPSC lines. For routine culture, mPSCs were maintained in flat bottom tissue culture plates (Falcon, Tewksbury, MA) coated with 0.2% gelatin. mPSCs were cultured using N2B27-based medium with addition of LIF and inhibitors for GSK3-beta and ERK (NB2iL) comprised of 48% Neurobasal Medium (Gibco, Gaithersburg, MD), 48% Dulbecco’s Modified Eagle Medium: Nutrient Mixture F-12 (Gibco, Gaithersburg, MD), 1% B-27 Plus Serum Free Supplements (Gibco, Gaithersburg, MD), 1% Glutamax, 1% 55mM 2-mercaptoethanol (Gibco, Gaithersburg, MD), 0.5% 100x N-2 Supplement (Gibco, Gaithersburg, MD), 0.5% Pen/Strep, 0.025% Bovine serum albumin (BSA, Wisent Bioproducts), 0.005% 200ng/ul LIF, 0.01% 30mM CHIR99021 (Tocris Bioscience), and 0.01% 10mM PD0325901 (STEMCELL Technologies). Medium was changed daily and mPSCs were passaged every 2 days (80% confluency) by exposing cells to Trypsin-EDTA (Gibco, Gaithersburg, MD) for 3 min at 37 degrees Celsius.

### Priming of mPSCs

To convert naive mPSC to a transient EpiLC state prior to differentiation, mPSCs were cultured in a priming medium (NBFAK) for two days before plating on micropatterns similar to what has been described previously in (Morgani et al. 2018). Briefly, naive mPSCs suspension were collected by applying 1ml/well of Trypsin-EDTA (Gibco, Gaithersburg, MD) to mPSCs grown in NB2iL medium and incubating them at 37°C for 3 minutes. 2mL of serum-containing medium was then added to neutralize trypsin activity. mPSCs were gently pipetted up and down to dissociate into a single cell suspension. Cells were collected at 400 rpm for 3 min and resuspended in NBFAK medium containing 48% Neurobasal Medium (Gibco, Gaithersburg, MD), 48% Dulbecco’s Modified Eagle Medium: Nutrient Mixture F-12 (Gibco, Gaithersburg, MD), 1% B-27 Plus Serum Free Supplements (Gibco, Gaithersburg, MD), 1% Glutamax, 1% 55mM 2-mercaptoethanol (Gibco, Gaithersburg, MD), 0.5% 100x N-2 Supplement (Gibco, Gaithersburg, MD), 0.5% Pen/Strep, 0.025% Bovine serum albumin (Wisent Bioproducts),12 ng/ml FGF2 (Peprotech, Rocky Hills, NJ), 20 ng/ml ACTIVIN A (Peprotech, Rocky Hills, NJ) and 1% KnockOut Serum Replacement (Gibco). Cells were counted and 200,000 cells were plated onto each well within a 6-well tissue culture plate (Falcon, Tewksbury, MA) coated with 0.2% gelatin. Cells were grown in NBFAK medium for 48 hours and medium was changed every day.

### Preparation of micropatterned plates

We followed our previously-developed method to create micropatterns using UV-lithography with glass slides and bottomless 96-well plates (Tewary et al. 2019). Briefly, glass cover-slips were cleaned with isopropanol and coated with a photo sensitive cell-phobic polymer using a spin coater at 2000rpm for 30s. 20 minutes of deep UV exposure was applied to the glass slide through a Quartz photomask with predefined micropatterns in a UV-Ozone cleaner (Jelight, Irvine, California) to photo-oxidize selected regions of the functionalized coating. Patterned glass slides were attached to bottomless 96-well plates using medical Epoxy (Henkel) with 4-hour cure time (in conventional oven at 54 degrees Celsius) to produce plates with patterned cell culture surfaces.

### Micropatterned mPSC colony plate preparation

Prior to seeding cells onto the plates, 100ul of ddH2O was added to each well followed by 25ul of 50mg/ml N-(3-Dimethylaminopropyl)-N-ethylcarbodiimide hydrochloride (Sigma, St Louis, Missouri) and 25ul of 50mg/ml N-Hydroxysuccinimide (Sigma, St Louis, Missouri) for 20 minutes to activate the wells. The wells were washed with ddH2O for 3 times and incubated with 100ul/well of 10μg/ml fibronectin bovine plasma (Sigma-Aldrich) in 0.1% gelatin for 2 hours on an orbital shaker at room temperature. The long incubation allowed for deposition and saturation of the extracellular matrix onto the functionalized regions of the glass. Wells were washed thrice with ddH2O prior to seeding to get rid of any passively absorbed ECM on non-functionalized regions. 50μL of ddH2O was pipetted into each well to ensure the ECM remains hydrated during preparation of the cell seeding suspension. ECM-coated plates were seeded with cells within 8 hours of assembly.

### Cell seeding and induction of gastrulation-like differentiation events in mPSC micropatterns

After 48-hour priming in NBFAK medium, single cell suspension of EpiLCs were collected by trypsinization as described above. Cells were centrifuged at 400 rpm for 5 mins and resuspended in NBFAK medium supplemented with 10 μM Rho-associated kinase inhibitor Y-27632 (ROCKi, Tocris Bioscience, UK). Cells were counted and seeded at 20 000 cells per well (or as described in the text) in 100ul/well of NBFAK with ROCKi. Plates were maintained in the tissue culture hood for 30 minutes after plating to allow time for cells to more evenly settle on the functionalized micropattern surface before moving to the incubator. After 24 hours, the medium was replaced with 200ul/well of fresh NBFAK medium (without ROCKi). After 48 hours upon cell seeding, the medium was changed to 200ul/well of NBFAWB medium containing 48% Neurobasal Medium, 48% Dulbecco’s Modified Eagle Medium: Nutrient Mixture F-12, 1% B-27 Plus Serum Free Supplements, 1% Glutamax, 1% 55mM 2-mercaptoethanol, 0.5% 100x N-2 Supplement, 0.5% Pen/Strep, 0.025% Bovine serum albumin,12 ng/ml FGF2, 20 ng/ml ACTIVIN A, 200ng/ml murine WNT3A (Peprotech, Rocky Hills, NJ), and 50ng/ml murine BMP4 (Peprotech, Rocky Hills, NJ). Cells were maintained in NBFAWB medium for up to 48 hours (or as described in the text) and medium was changed daily.

### Fixing and immunofluorescence staining of differentiated mPSCs on micropatterns

Staining followed a protocol previously developed to stain tumor spheres (Weiswald et al. 2010) to ensure sufficient penetration even in thick colonies. Micropatterned colonies were fixed with 4% paraformaldehyde in PBS containing 1% Triton-X (Sigma) (PBST) for 2 hours at 4°C. Cells were then washed thrice with PBS and blocked in 0.1% PBST supplemented with 2% BSA (Wisent Bioproducts) for 30 minutes at room temperature. Cells were then immunostained overnight at 4°C with primary antibodies (SOX2 (1:1200, Invitrogen #14-9811-82) and BRA (1:400, R&D #AF2085)) diluted in 0.1% PBST. Cells were washed thrice with 0.1% PBST on the following day, and were then immunostained overnight at 4°C with secondary antibodies (1:200, Donkey anti Goat IgG (H+L) Secondary Antibody, Alexa Fluor 647, Invitrogen; 1:200, Donkey anti Rat IgG (H+L) Highly Cross Adsorbed Secondary Antibody, Alexa Fluor 488, Invitrogen) and DAPI (1:1000, Sigma #D9542) diluted in 0.1% PBST. On the following day, cells were washed thrice with 0.1% PBST and stored at 4°C with PBS. Micropatterned plates were imaged using an automated imaging pipeline on the Zeiss LSM800 confocal with a 20x 0.8 NA air objective acquiring five z-slices per imaged field, and imaging around 90% of each well. Images were reduced the maximum intensity projects before analysis.

### Sample definition and in-laboratory replication

A minimum of two independent experiments were done for each experiment, with three technical replicates for each experiment.

### Image quantification

To be able to analyze the wealth of data generated from our micropatterned high-throughput platform in a systematic manner, we developed an analytic pipeline tailored for this purpose. The design goals of this platform were to build a semi-automated approach to quantify cell fate organization in colonies of cells, and to create this software framework in an environment that is openly accessible and easy to use without significant computational training while still allowing great flexibility and power for experienced users. While this pipeline was initially intended for general use, the user-interfacing parts are not yet completely implemented so in this study we relied on using parts of the pipeline programmatically and making modifications to the code as needed.

We implemented our framework as a set of graphical widgets in an open programmable environment. Specifically, we used Jupyter Notebooks (Kluyver et al. 2016; Pérez and Granger 2007) via JupyterLab, as they allow for display of rich content inline with code in the same browser-based interface. This allowed us to leverage web technologies such as HTML and Javascript to create interactive widgets via Panel (Rudiger et al. 2020) and Param (Stevens et al. 2020), and interactive plots using Bokeh (Bokeh Development Team 2020) and Holoviews (Rudiger et al. 2020). Interactivity is a central component of our analytical framework as widgets makes it widely accessible to a large number of scientists and speeds up parameter optimization as users can see changes represented in the microscopy images in real time when they drag sliders or in other ways update the widgets. The inclusion of interactive plots provides rich information on individual data points and through the display of images for selected data points; this allows us to directly relate measurements back to microscopy images. This also contributes to the ease of use of the platform since for every data point it is easy to view the image of the cell colony, which simplifies the intuitive understanding of how metrics are derived and reduces the risk of introducing unseen errors.

Our analytical framework relies on several well-established scientific Python packages to perform the quantification of image data. Specifically, we use pandas (Reback et al. 2020; Wes McKinney 2010) as the backbone for storing and operating on data using dataframes, scikit-image (van der Walt et al. 2014) and numpy (van der Walt, Colbert, and Varoquaux 2011) for image processing, statsmodels (Seabold and Perktold 2010), scipy (Jones et al. 2001), scikit-learn (Pedregosa et al. 2011), and scikits-bootstrap for deriving statistical quantities of data, joblib (Joblib Development Team 2020) and pandarallel to increase performance through parallel computing, and numba (Lam, Pitrou, and Seibert 2015) to further speed up computations by compiling custom Python functions at runtime.

There are three main steps involves in the analysis process, first colonies of cells are identified in microscopy images, which requires the use of a nuclear dye to stain all cells in the colony. Identified colonies can be filtered based on metrics such as heir area, circularity, solidity (area ratio to their bounding box), and proximity to the image border. To identify suitable thresholds for these parameters, we use scatter plots similar to how subpopulations are identified using gating in flow cytometry. Since it is not immediately obvious where a certain threshold should lie just from the data points in the plot, selection of data points displays the corresponding microscopy images to facilitate the decision whether these data points should be included or excluded from the analysis.

Details for how to use the image quantification pipeline can be found in the code repository at https://gitlab.com/stemcellbioengineering/colony-image-analysis. Briefly, there are three main steps involved in the analysis process. First cells are classified into colonies based on a nuclear stain and filtered based on their area, circularity, solidity, and proximity to the image border. Second, a threshold is set for which pixels are classified as positive and which are negative using a semi-automatic approach based on the shape of the pixel intensity curve. Third, colony metrics are calculated, including shape properties such as area, aspect ratio (elongation), solidity, and circularity which allows us to stratify colonies by shape in assays where this is of interest. The center of mass is calculated for positive pixel regions and the (diameter-normalized) distance between the center of mass for different markers can then be used to determine its offset from the colony centroid, which is a measure of the degree of asymmetrical expression in the colony. To easily compare distributions of measurements between different conditions, we developed a variation strip/scatter plots where points are laid out according to their kernel density estimate. The shape of this distribution is analogous to a violin plot, but does not have the drawbacks of masking the number of observations, and making small distributions appear unnaturally smooth.

### Solution of the activator equation in steady state

To simplify the calculations for the steady-state solution of the concentration of the activator. We first consider that parameter h in the source term is a very high number. This permits to simplify the source term from *P(A,B)=A*^*h*^*/(A*^*h*^*+B*^*h*^) to a step function where *P(A,B)=*1 for *A>B* and *P(A,B)*=0 otherwise. Then, the general solution for **Eq. 5** with boundary conditions as in **Eq. 7** reads:

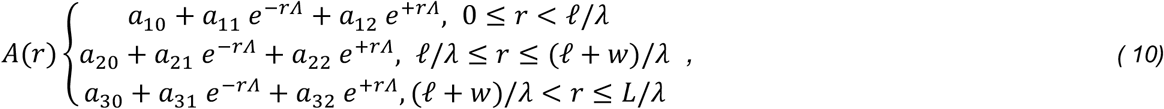

with *w* the width of the source region of the activation *A*, and *ℓ* the distance from the center of the simulated colony to the source region *w*.

The expression for the constants *a*_*ij*_ read:

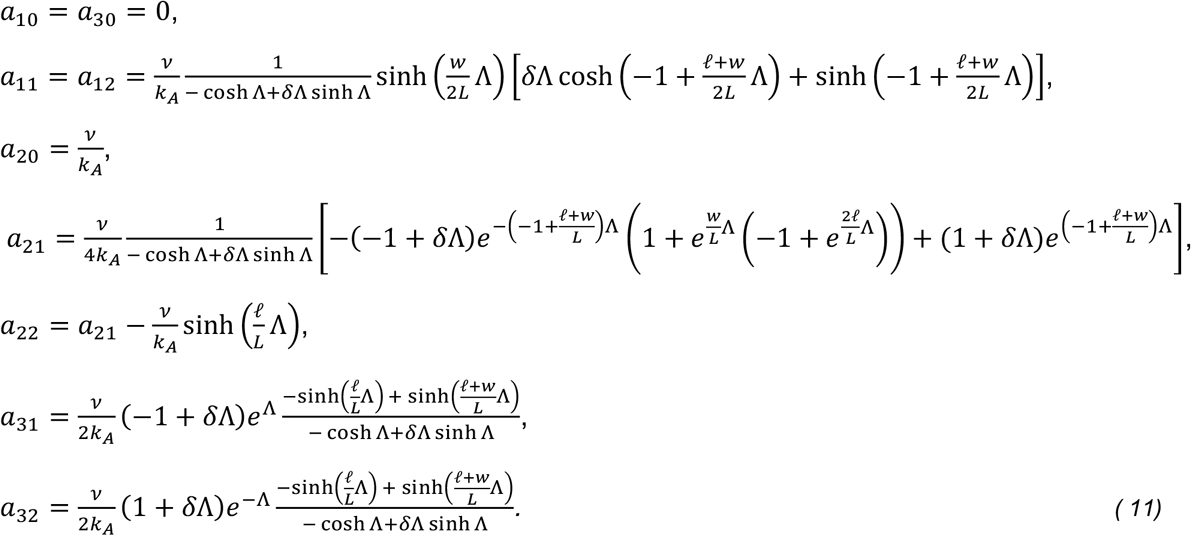

### Model fit to BRA intensity gradients

To calculate the length-scale of the BRA gradients, we use the analytical steady-state solutions of our mathematical model in a non-dimensional form (see sup. Mat.) to fit the experimental data. The solutions of our model indicate that we should expect a decay in the intensity profile that aligns with a function that combines exponential terms, as in **Eq. 9**, more precisely the fit function follows the form *a* + *b* cosh*A*(*c r*), with constant parameters *a,b* and *c* as corresponds to the branch for 0 ≤ *r* < *ℓ*/*λ* in **Eq. 10**, considering that *a*≠0.

The experimental gradients were extracted using Context Explorer (Ostblom et al. 2019), and as fitting routine we used Matlab tool *cftool* to quantify each profile.

### Simulations in circular geometries

Simulations are done using Comsol Multiphysics 5.6.

## Acknowledgements

We want to thank the entire Stem Cell Bioengineering (Zandstra) Lab for fruitful discussions and critical feedback. We acknowledge funding support from the following grants: NSERC Discovery Grant – number RGPIN-2020-06496, CIHR Foundation Grant – number FDN 154283, Canada Foundation for Innovation - John R. Evans Leaders Fund - Project number 37472; and Innovation Fund - Project number 36275.

---

**Supplementary Figure S1:**
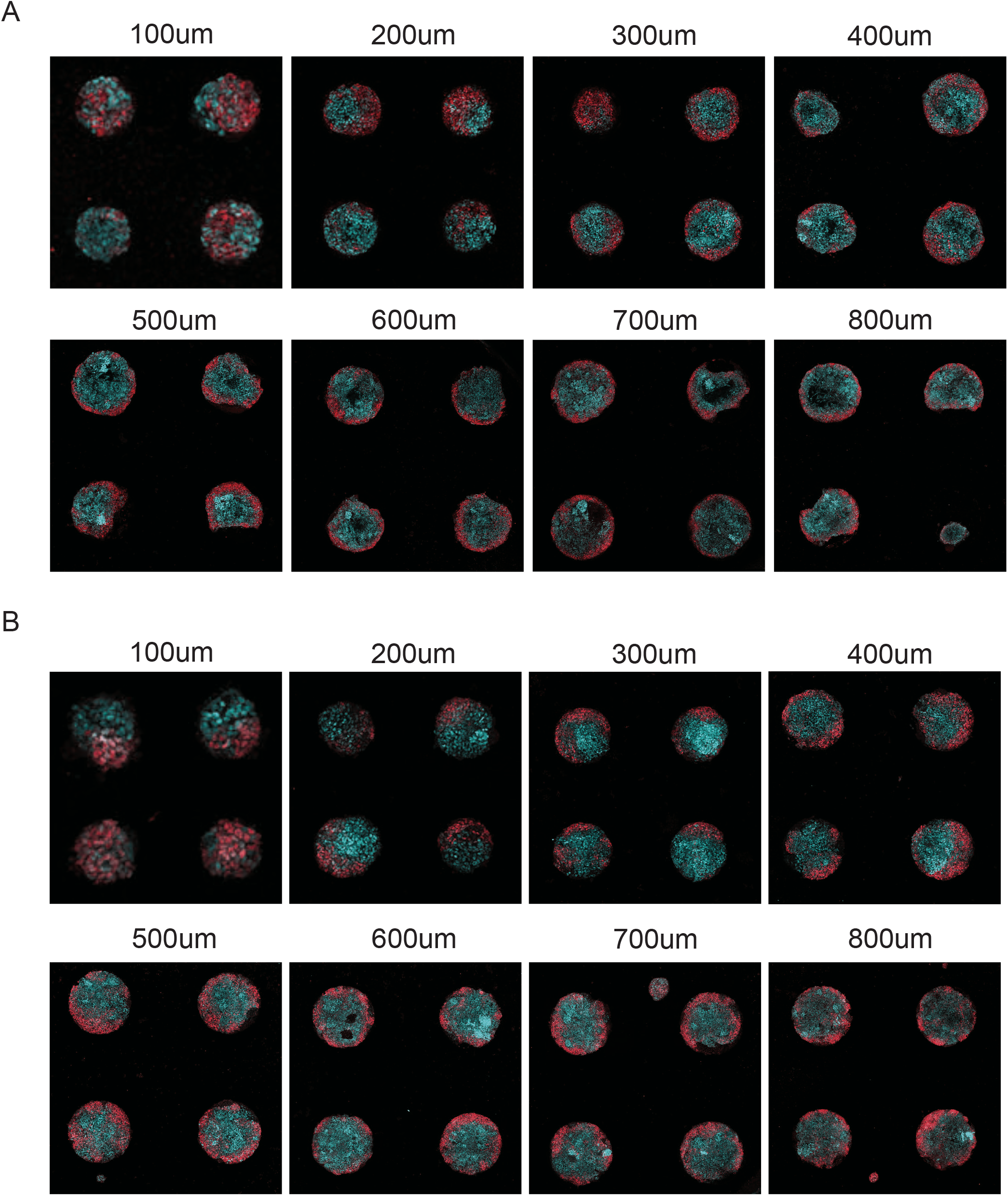
A critical range of colony diameters supports polarized expression of Bra and Sox2. **A**,**B)** Representative images of **(A)** R1 and **(B)** BraGFP/+ mPSC micropatterned colonies of 100μm to 800μm diameter after 48hr differentiation. Cyan = Sox2, Red = Bra.

**Supplementary Figure S2:**
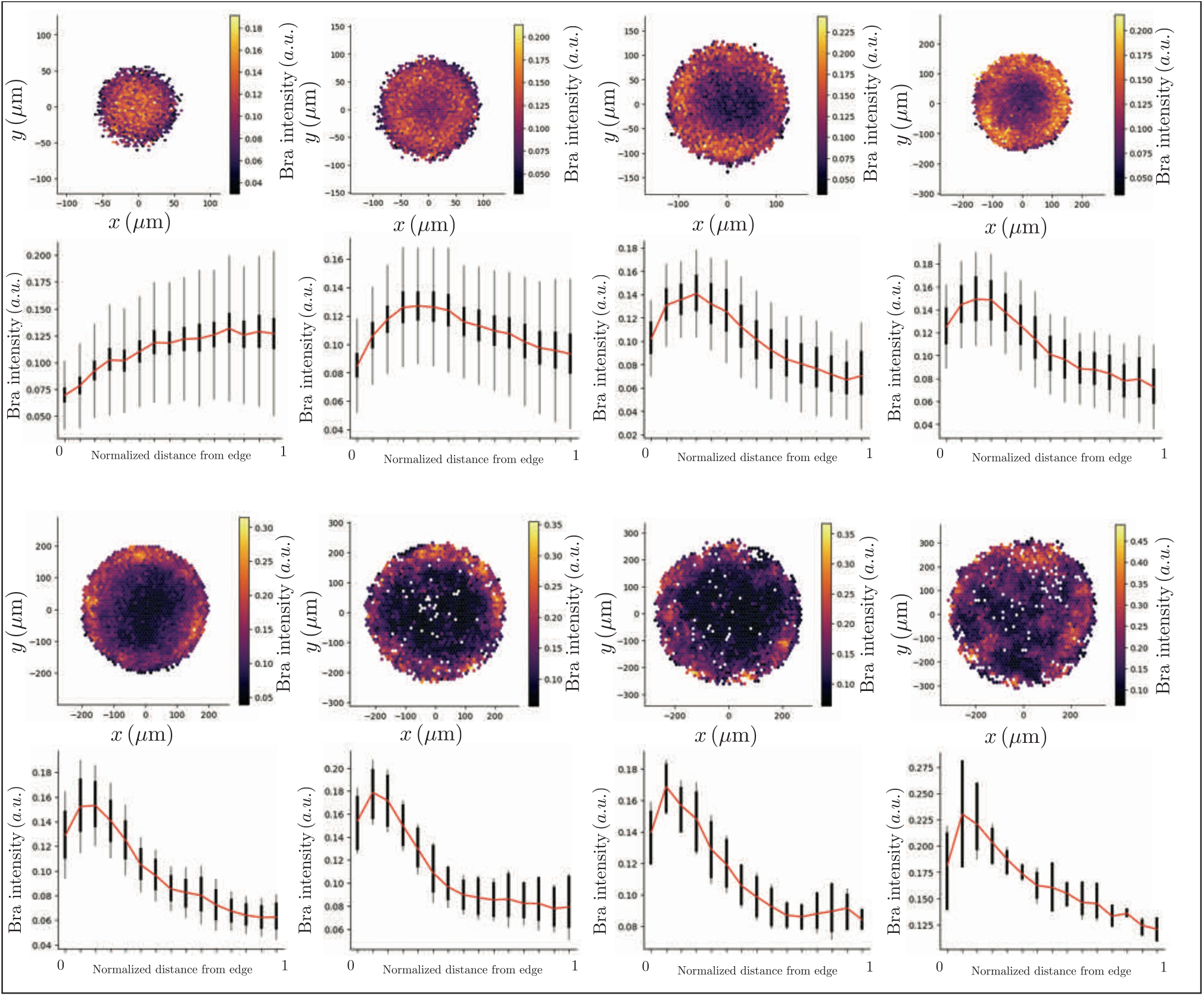
Quantification of average BRA expression gradients (seeding cell density 10k cells/well). Heatmaps showing the average Bra expression over a total of N=307. From upper-left corner to bottom right, the radius of the colony spans L=50-400μm. Below each heatmap, a radial quantification of Bra gradient shown in bins over a normalized radius. Red line connects the average value per bin. Thick back lines represent the 95% confidence interval for cells grouped in that bin, and light black lines the standard deviation. Cell seeding density 10k cells per well.

**Supplementary Figure S3:**
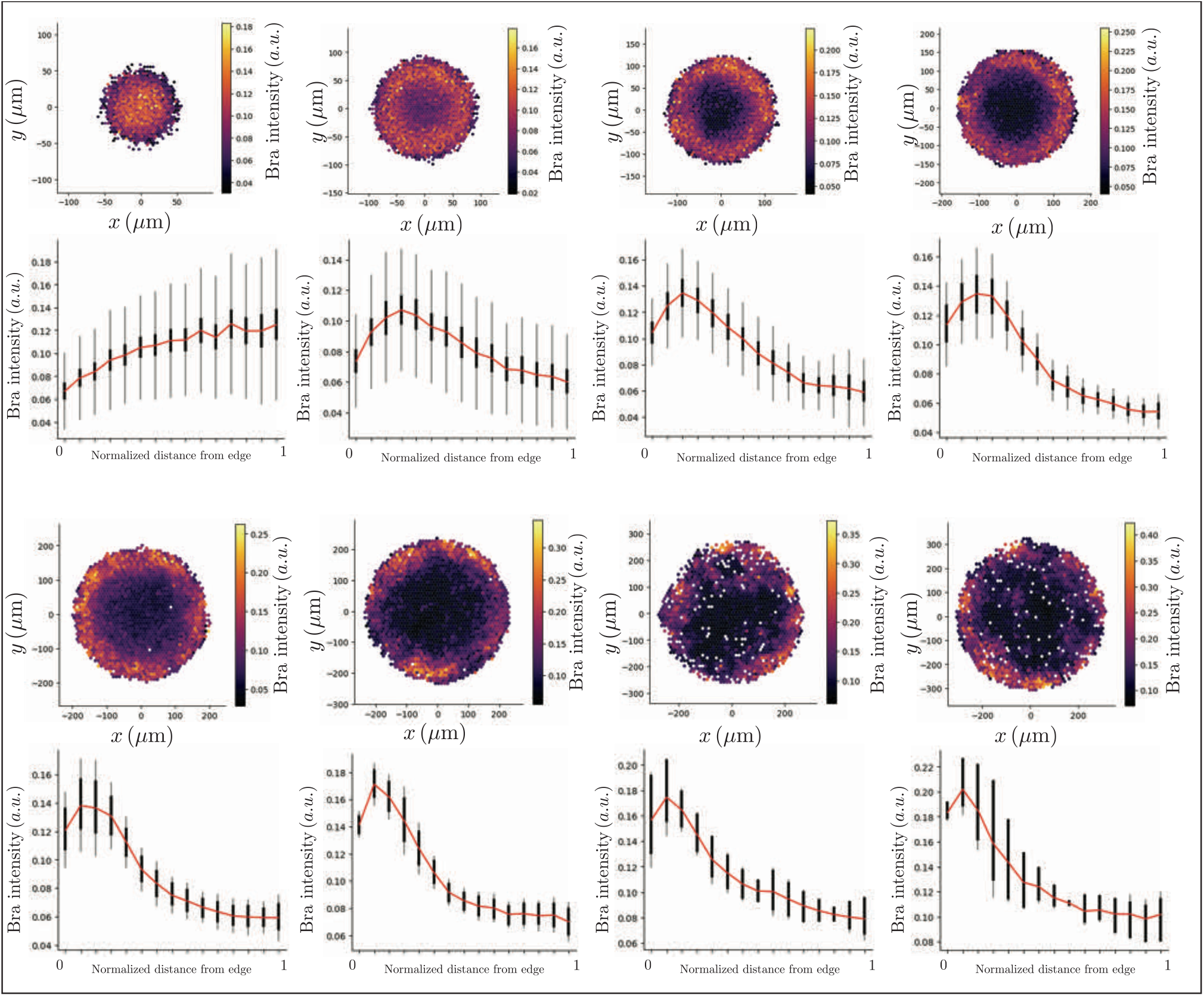
Quantification of average BRA expression gradients (seeding cell density 12k cells/well). Heatmaps showing the average Bra expression over a total of N=303. From upper-left corner to bottom right, the radius of the colony spans L=50-400μm. Below each heatmap, a radial quantification of Bra gradient shown in bins over a normalized radius. Red line connects the average value per bin. Thick back lines represent the 95% confidence interval for cells grouped in that bin, and light black lines the standard deviation. Cell seeding density 12k cells per well.

**Supplementary Figure S4:**
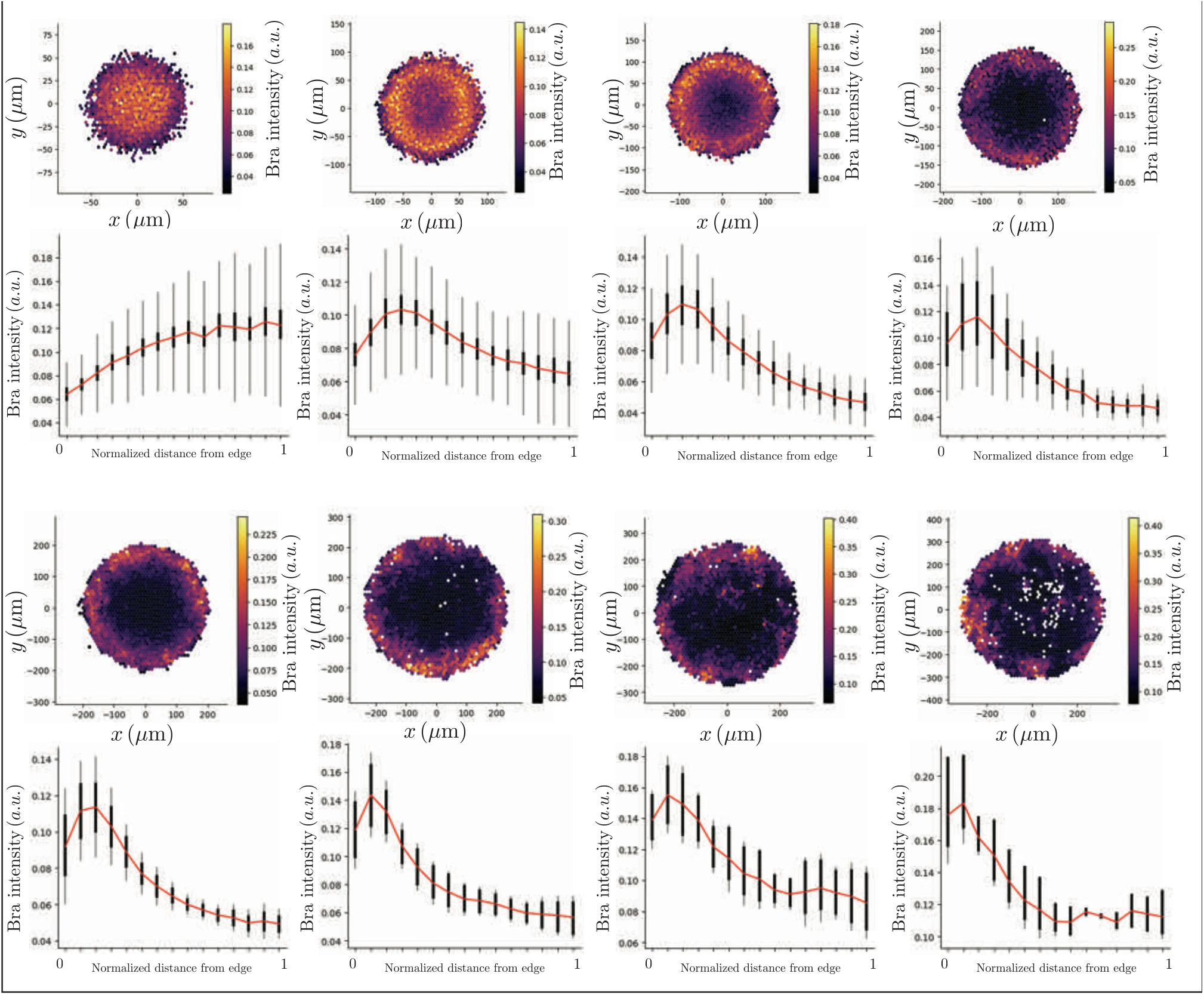
Quantification of average BRA expression gradients (seeding cell density 14k cells/well). Heatmaps showing the average Bra expression over a total of N=306. From upper-left corner to bottom right, the radius of the colony spans L=50-400μm. Below each heatmap, a radial quantification of Bra gradient shown in bins over a normalized radius. Red line connects the average value per bin. Thick back lines represent the 95% confidence interval for cells grouped in that bin, and light black lines the standard deviation. Cell seeding density 14k cells per well.

## Notes

### Competing Interest Statement

The authors have declared no competing interest.

## References

Aguilar-Hidalgo, Daniel, Zena Hadjivasilou, Maria Romanova-Michaelides, Marcos González-Gaitán, and Frank Jülicher. 2019. “Dynamic Modes of Morphogen Transport.” https://doi.org/10.48550/ARXIV.1909.13280.

Aguilar-Hidalgo, Daniel, Steffen Werner, Ortrud Wartlick, Marcos González-Gaitán, Benjamin M. Friedrich, and Frank Jülicher. 2018. “Critical Point in Self-Organized Tissue Growth.” Physical Review Letters 120 (19): 198102. https://doi.org/10.1103/PhysRevLett.120.198102.

Amadei, Gianluca, Charlotte E. Handford, Chengxiang Qiu, Joachim De Jonghe, Hannah Greenfeld, Martin Tran, Beth K. Martin, et al. 2022. “Embryo Model Completes Gastrulation to Neurulation and Organogenesis.” Nature 610 (7930): 143–53. https://doi.org/10.1038/s41586-022-05246-3.

Beccari, Leonardo, Naomi Moris, Mehmet Girgin, David A. Turner, Peter Baillie-Johnson, Anne-Catherine Cossy, Matthias P. Lutolf, Denis Duboule, and Alfonso Martinez Arias. 2018. “Multi-Axial Self-Organization Properties of Mouse Embryonic Stem Cells into Gastruloids.” Nature. Nature Publishing Group. https://doi.org/10.1038/s41586-018-0578-0.

Bernardo, Andreia S., Tiago Faial, Lucy Gardner, Kathy K. Niakan, Daniel Ortmann, Claire E. Senner, Elizabeth M. Callery, et al. 2011. “BRACHYURY and CDX2 Mediate BMP-Induced Differentiation of Human and Mouse Pluripotent Stem Cells into Embryonic and Extraembryonic Lineages.” Cell Stem Cell 9 (2): 144–55. https://doi.org/10.1016/j.stem.2011.06.015.

Brassard, Jonathan A., and Matthias P. Lutolf. 2019. “Engineering Stem Cell Self-Organization to Build Better Organoids.” Cell Stem Cell 24 (6): 860–76. https://doi.org/10.1016/j.stem.2019.05.005.

Chhabra, Sapna, Lizhong Liu, Ryan Goh, Xiangyu Kong, and Aryeh Warmflash. 2019. “Dissecting the Dynamics of Signaling Events in the BMP, WNT, and NODAL Cascade during Self-Organized Fate Patterning in Human Gastruloids.” PLOS Biology 17 (10): e3000498. https://doi.org/10.1371/journal.pbio.3000498.

Chio, Chung Chi, and Ying-Lung Steve Tse. 2020. “Hindered Diffusion near Fluid–Solid Interfaces: Comparison of Molecular Dynamics to Continuum Hydrodynamics.” Langmuir 36 (32): 9412–23. https://doi.org/10.1021/acs.langmuir.0c01228.

Dillon, R., P. K. Maini, and H. G. Othmer. 1994. “Pattern Formation in Generalized Turing Systems: I. Steady-State Patterns in Systems with Mixed Boundary Conditions.” Journal of Mathematical Biology 32 (4): 345–93. https://doi.org/10.1007/BF00160165.

Eloul, Shaltiel, and Richard G. Compton. 2016. “General Model of Hindered Diffusion.” The Journal of Physical Chemistry Letters 7 (21): 4317–21. https://doi.org/10.1021/acs.jpclett.6b02275.

Erban, Radek, and S Jonathan Chapman. 2007. “Reactive Boundary Conditions for Stochastic Simulations of Reaction–Diffusion Processes.” Physical Biology 4 (1): 16–28. https://doi.org/10.1088/1478-3975/4/1/003.

Etoc, Fred, Jakob Metzger, Albert Ruzo, Christoph Kirst, Anna Yoney, M. Zeeshan Ozair, Ali H. Brivanlou, and Eric D. Siggia. 2016. “A Balance between Secreted Inhibitors and Edge Sensing Controls Gastruloid Self-Organization.” Developmental Cell 39 (3): 302–15. https://doi.org/10.1016/j.devcel.2016.09.016.

Fattah, Abdel Rahman Abdel, Sergei Grebenyuk, Idris Salmon, and Adrian Ranga. 2021. “Neuroepithelial Organoid Patterning Is Mediated by Wnt-Driven Turing Mechanism.” Preprint. Developmental Biology. https://doi.org/10.1101/2021.01.11.426254.

Gjorevski, N., M. Nikolaev, T. E. Brown, O. Mitrofanova, N. Brandenberg, F. W. DelRio, F. M. Yavitt, P. Liberali, K. S. Anseth, and M. P. Lutolf. 2022. “Tissue Geometry Drives Deterministic Organoid Patterning.” Science 375 (6576): eaaw9021. https://doi.org/10.1126/science.aaw9021.

Harrison, Sarah Ellys, Berna Sozen, Neophytos Christodoulou, Christos Kyprianou, and Magdalena Zernicka-Goetz. 2017. “Assembly of Embryonic and Extraembryonic Stem Cells to Mimic Embryogenesis in Vitro.” Science 356 (6334): eaal1810. https://doi.org/10.1126/science.aal1810.

Ishihara, Keisuke, and Elly M. Tanaka. 2018. “Spontaneous Symmetry Breaking and Pattern Formation of Organoids.” Current Opinion in Systems Biology 11 (October): 123–28. https://doi.org/10.1016/j.coisb.2018.06.002.

Manfrin, Andrea, Yoji Tabata, Eric R. Paquet, Ambroise R. Vuaridel, François R. Rivest, Felix Naef, and Matthias P. Lutolf. 2019. “Engineered Signaling Centers for the Spatially Controlled Patterning of Human Pluripotent Stem Cells.” Nature Methods 16 (7): 640–48. https://doi.org/10.1038/s41592-019-0455-2.

Morgani, Sophie M, Jakob J Metzger, Jennifer Nichols, Eric D Siggia, and Anna-Katerina Hadjantonakis. 2018. “Micropattern Differentiation of Mouse Pluripotent Stem Cells Recapitulates Embryo Regionalized Cell Fate Patterning.” ELife 7 (February): e32839. https://doi.org/10.7554/eLife.32839.

Moris, Naomi, Kerim Anlas, Susanne C. van den Brink, Anna Alemany, Julia Schröder, Sabitri Ghimire, Tina Balayo, Alexander van Oudenaarden, and Alfonso Martinez Arias. 2020. “An in Vitro Model of Early Anteroposterior Organization during Human Development.” Nature 582 (7812): 410–15. https://doi.org/10.1038/s41586-020-2383-9.

Müller, Patrick, Katherine W. Rogers, Shuizi R. Yu, Michael Brand, and Alexander F. Schier. 2013. “Morphogen Transport.” Development 140 (8): 1621–38. https://doi.org/10.1242/dev.083519.

Muncie, Jonathon M., Nadia M.E. Ayad, Johnathon N. Lakins, Xufeng Xue, Jianping Fu, and Valerie M. Weaver. 2020. “Mechanical Tension Promotes Formation of Gastrulation-like Nodes and Patterns Mesoderm Specification in Human Embryonic Stem Cells.” Developmental Cell 55 (6): 679-694.e11. https://doi.org/10.1016/j.devcel.2020.10.015.

Nemashkalo, Anastasiia, Albert Ruzo, Idse Heemskerk, and Aryeh Warmflash. 2017. “Morphogen and Community Effects Determine Cell Fates in Response to BMP4 Signaling in Human Embryonic Stem Cells.” Development 144 (17): 3042–53. https://doi.org/10.1242/dev.153239.

Orietti, Lorenzo C., Viviane Souza Rosa, Francesco Antonica, Christos Kyprianou, William Mansfield, Henrique Marques-Souza, Marta N. Shahbazi, and Magdalena Zernicka-Goetz. 2021. “Embryo Size Regulates the Timing and Mechanism of Pluripotent Tissue Morphogenesis.” Stem Cell Reports 16 (5): 1182–96. https://doi.org/10.1016/j.stemcr.2020.09.004.

Ostblom, Joel, Emanuel J. P. Nazareth, Mukul Tewary, and Peter W. Zandstra. 2019. “Context-Explorer: Analysis of Spatially Organized Protein Expression in High-Throughput Screens.” PLOS Computational Biology 15 (1): e1006384. https://doi.org/10.1371/journal.pcbi.1006384.

Romanova-Michaelides, Maria, Zena Hadjivasiliou, Daniel Aguilar-Hidalgo, Dimitris Basagiannis, Carole Seum, Marine Dubois, Frank Jülicher, and Marcos Gonzalez-Gaitan. 2022. “Morphogen Gradient Scaling by Recycling of Intracellular Dpp.” Nature 602 (7896): 287–93. https://doi.org/10.1038/s41586-021-04346-w.

Sozen, Berna, Jake Cornwall-Scoones, and Magdalena Zernicka-Goetz. 2021. “The Dynamics of Morphogenesis in Stem Cell-Based Embryology: Novel Insights for Symmetry Breaking.” Developmental Biology 474 (June): 82–90. https://doi.org/10.1016/j.ydbio.2020.12.005.

Tewary, Mukul, Dominika Dziedzicka, Joel Ostblom, Laura Prochazka, Nika Shakiba, Tiam Heydari, Daniel Aguilar-Hidalgo, et al. 2019. “High-Throughput Micropatterning Platform Reveals Nodal-Dependent Bisection of Peri-Gastrulation– Associated versus Preneurulation-Associated Fate Patterning.” PLOS Biology 17 (10): e3000081. https://doi.org/10.1371/journal.pbio.3000081.

Tewary, Mukul, Joel Ostblom, Laura Prochazka, Teresa Zulueta-Coarasa, Nika Shakiba, Rodrigo Fernandez-Gonzalez, and Peter W. Zandstra. 2017. “A Stepwise Model of Reaction-Diffusion and Positional Information Governs Self-Organized Human Peri-Gastrulation-like Patterning.” Development 144 (23): 4298–4312. https://doi.org/10.1242/dev.149658.

Turing, A. M. 1952. “The Chemical Basis of Morphogenesis.” Bulletin of Mathematical Biology 52 (1): 153–97. https://doi.org/10.1007/BF02459572.

Turner, David A., Mehmet Girgin, Luz Alonso-Crisostomo, Vikas Trivedi, Peter Baillie-Johnson, Cherise R. Glodowski, Penelope C. Hayward, et al. 2017. “Anteroposterior Polarity and Elongation in the Absence of Extra-Embryonic Tissues and of Spatially Localised Signalling in Gastruloids: Mammalian Embryonic Organoids.” Development 144 (21): 3894–3906. https://doi.org/10.1242/dev.150391.

Warmflash, Aryeh, Benoit Sorre, Fred Etoc, Eric D Siggia, and Ali H Brivanlou. 2014. “A Method to Recapitulate Early Embryonic Spatial Patterning in Human Embryonic Stem Cells.” Nature Methods 11 (8): 847–54. https://doi.org/10.1038/nmeth.3016.

Werner, Steffen, Tom Stückemann, Manuel Beirán Amigo, Jochen C. Rink, Frank Jülicher, and Benjamin M. Friedrich. 2015. “Scaling and Regeneration of Self-Organized Patterns.” Physical Review Letters 114 (13): 138101. https://doi.org/10.1103/PhysRevLett.114.138101.

